# Mag-Net: Rapid enrichment of membrane-bound particles enables high coverage quantitative analysis of the plasma proteome

**DOI:** 10.1101/2023.06.10.544439

**Authors:** Christine C. Wu, Kristine A. Tsantilas, Jea Park, Deanna Plubell, Justin A. Sanders, Previn Naicker, Ireshyn Govender, Sindisiwe Buthelezi, Stoyan Stoychev, Justin Jordaan, Gennifer Merrihew, Eric Huang, Edward D. Parker, Michael Riffle, Andrew N. Hoofnagle, William S. Noble, Kathleen L. Poston, Thomas J. Montine, Michael J. MacCoss

## Abstract

Membrane-bound particles in plasma are composed of exosomes, microvesicles, and apoptotic bodies and represent ∼1-2% of the total protein composition. Proteomic interrogation of this subset of plasma proteins augments the representation of tissue-specific proteins, representing a “liquid biopsy,” while enabling the detection of proteins that would otherwise be beyond the dynamic range of liquid chromatography-tandem mass spectrometry of unfractionated plasma. We have developed an enrichment strategy (Mag-Net) using hyper-porous strong-anion exchange magnetic microparticles to sieve membrane-bound particles from plasma. The Mag-Net method is robust, reproducible, inexpensive, and requires <100 μL plasma input. Coupled to a quantitative data-independent mass spectrometry analytical strategy, we demonstrate that we can collect results for >37,000 peptides from >4,000 plasma proteins with high precision. Using this analytical pipeline on a small cohort of patients with neurodegenerative disease and healthy age-matched controls, we discovered 204 proteins that differentiate (q-value < 0.05) patients with Alzheimer’s disease dementia (ADD) from those without ADD. Our method also discovered 310 proteins that were different between Parkinson’s disease and those with either ADD or healthy cognitively normal individuals. Using machine learning we were able to distinguish between ADD and not ADD with a mean ROC AUC = 0.98 ± 0.06.

## INTRODUCTION

The robust quantitative characterization of proteins from plasma is critical to the diagnosis of disease and monitoring progression as well as response to treatment. Blood plasma is the primary specimen type analyzed in clinical laboratories, and proteins in plasma hold exceptional promise as they facilitate the ubiquitous sampling of tissues while remaining minimally invasive and, in essence, providing a proxy “liquid biopsy” of tissues. Despite blood plasma being estimated to contain well over 10,000 proteins, most proteomics experiments performed using unfractionated plasma digests are limited to measuring <800 proteins^1–4^, mainly due to the immense dynamic range of protein concentrations spanning 10 orders of magnitude -- e.g., the difference in concentration between albumin and IL-6^5^. In other words, 50% of plasma protein by mass is one protein (albumin), and 22 proteins account for 99% of the total protein mass^6,7^.

The dynamic range of plasma proteins can be partially mitigated by depleting the most abundant proteins using immunoaffinity subtraction chromatography. Such depletion frequently removes 6-20 of the most abundant proteins in human plasma by about 90-99%^8–10^. Depletion increases the number of detected proteins by decreasing the dynamic range. However, these depletion columns also capture more than just the target antigens, frequently removing the target antigens plus any bound proteins. Finally, even after immunodepleting the most abundant proteins, the 50 most abundant plasma proteins can still account for 90% of the total signal in a proteomics experiment^10^.

To further overcome the dynamic range challenge, protein depletion can be combined with off-line protein or peptide level fractionation before LC-MS/MS. There have been numerous strategies, including SDS-PAGE, isoelectric focusing, peptide level high-pH reversed-phase fractionation, and peptide level strong cation exchange separations. Downsides of these methods are that they have the disadvantages of immunodepletion and also result in a major reduction in throughput – some methods resulting in as many as 25-96 fractions per sample^11–13^.

Recently, Tognetti et al demonstrated the value of state-of-the-art data independent acquisition methods – both acquisition methods and computational analysis – for plasma proteomics^14^. By combining Agilent affinity removal column human-14, with the acquisition of DIA with a 2-hour HPLC gradient on an Orbitrap Exploris 480 mass spectrometer (Thermo Fisher Scientific), they could measure 2732 proteins across their samples without additional fractionation. This work is significant because it was the first to demonstrate that >2500 proteins could be measured in plasma across multiple samples using a single LC-MS/MS run per sample.

More recently, Blume et al. reported the enrichment of different subpopulations of the plasma proteome without immunoaffinity depletion using paramagnetic nanoparticles derivatized with various surface chemistries^15^. These data show that using multiple nanoparticles and LC-MS runs can detect 1500 to 2000 protein groups^15–17^. However, like prior fractionation experiments, the number of nanoparticles used for enrichment comes at the expense of throughput. Despite these challenges, the excitement and current impact of this nanoparticle technology highlights the promise in using a magnetic bead to capture a subset of the plasma proteome to extend the analysis to lower abundant proteins.

Plasma contains a heterogeneous collection of membrane-bound particles, commonly referred to as extracellular vesicles (EV), which include exosomes, microvesicles, and apoptotic bodies. These membrane-bound particles represent ∼1-2% of the total protein composition in plasma and are commonly enriched using a variety of methods, such as ultracentrifugation, size exclusion chromatography, and immunoaffinity capture. Kverneland et al. recently demonstrated that by using an EV fraction prepared by differential ultracentrifugation coupled with the latest DIA-mass spectrometry methods, they could measure almost 4000 proteins in a single LC-MS/MS run without depletion^18^. Analysis of this enriched fraction by mass spectrometry may provide deeper insight into cell type-specific processes that are generating these different types of EV and potentially improve the range of the proteins measurable in plasma. However, EV isolation by ultracentrifugation is time consuming and requires specialized centrifuge equipment capable of performing at >100,000 x g, making this technology difficult to apply to large clinical cohorts. To address this limitation, Kverneland et al. used multiple ultracentrifugation steps at 20,000 x g to collect a pellet that is enriched in larger >100 nm diameter membrane-based particles^18^. These data demonstrate the potential of EVs as useful subfraction of the plasma proteome that improves the coverage of biomedically interesting proteins. However, EV enrichment protocols based entirely on density are often contaminated with high density lipoprotein (HDL) particles as they have similar density^19^. These approaches demonstrate the need for an EV enrichment protocol that can minimize contamination with lipoprotein particles and can be performed in a 96-well plate format.

A magnetic bead-based chemical affinity purification strategy for the enrichment of EVs from biofluids was recently described by Tao and colleagues^20–22^. This “EVTrap” strategy uses a combination of “hydrophilic and aromatic lipophilic groups” that the authors describe as having high affinity toward lipid-coated EVs^20,21^. The reported methodology is promising and demonstrated significant membrane particle enrichment with minimal contamination relative to competing methods like ultracentrifugation, size exclusion, and polymer precipitation^20^. Furthermore, the use of magnetic beads to capture EVs makes this strategy amendable to automation. Using this simple magnetic bead capture method, they reported the measurement of >1,000 proteins from just 10 µl of plasma and 600-1,000 phosphoproteins from 0.5 mL of plasma^21^. This method has even been used to define putative diagnostic biosignatures from urinary EVs for Parkinson’s disease^22^. Although promising, the EVTrap magnetic beads are not currently commercially available. Thus, the scientific community remains in need of a strategy for the enrichment of EV particles that is broadly accessible to enable assay transferability and harmonization within the community.

Here we describe Mag-Net, a simple, inexpensive, and robust magnetic bead-based method to capture EV particles from plasma while simultaneously depleting abundant plasma proteins. The method enriches EVs from plasma using a combination of size and charge. Coupled to a quantitative data-independent mass spectrometry analytical strategy, we demonstrate routine measurement of >37,000 peptides from >4,000 plasma proteins with high precision. Interestingly, 21.4% proteins measured by Mag-Net have a predicted transmembrane domain by either Philius^23^ or DeepTMHMM^24^, with an additional 5-10% known to be lipid-anchored or peripherally associated membrane proteins, a category of proteins often under-represented in proteomic analyses, particularly from plasma. We characterize the proteins captured by Mag-Net, the physical characteristics of the captured membrane particles, and demonstrate the value of this method for high coverage quantitative analysis of the plasma proteome. To illustrate the significance of this pipeline, we analyzed plasma samples collected from a small cohort of 40 individuals: 10 with Alzheimer’s disease dementia (ADD); 10 Parkinson’s disease (PD) cognitively normal (PDCN); 10 PD with dementia (PDD); and 10 age-matched healthy and cognitively normal (HCN) individuals. Our findings demonstrate the ability to distinguish between individuals with and without dementia, regardless of the cause (i.e. Alzheimer’s or Parkinson’s disease). Strikingly, also we discovered proteins that can distinguish between individuals with cognitive impairment caused by different neurodegenerative diseases.

## RESULTS AND DISCUSSION

### Overview of the Mag-Net method for plasma proteomics

To improve the dynamic range of plasma proteomics, we endeavored to develop a simple, low-cost, efficient, and automated method capable of enriching membrane-bound particles while simultaneously depleting abundant plasma proteins. MagReSyn® magnetic particles have been demonstrated to have high binding capacity and are compatible with automated LC-MS/MS workflows^25–27^. The MagReSyn® beads have a hyper-porous polymer matrix that enables biomolecules to penetrate throughout the volume of the particles. Previously, positively charged peptides composed of polylysine (K8, K16) have been shown to bind phospholipids and enable the enrichment of extracellular vesicles^28^. Therefore, using an electrostatic strategy, we tested the MagReSyn® strong anion exchange (SAX) beads, functionalized with quaternary ammonium, for binding to phospholipid bilayer-bound particles and depletion of abundant plasma proteins in plasma. Figure 1 illustrates the Mag-Net workflow. Plasma is incubated with SAX beads in a 4:1 volume ratio and incubated at room temperature with gentle mixing. Unbound plasma components are washed away and depleted. The captured membrane particles are lysed with SDS, and the liberated proteins are reduced, alkylated, and prepared using protein aggregation capture (PAC) sample preparation^25,29^. The proteins were aggregated to the SAX beads using acetonitrile, washed with multiple cycles of acetonitrile and ethanol, and then digested with trypsin. The entire process was performed using a KingFisher Flex magnetic bead robot using two separate cycles as illustrated in Figure 1b.

**Figure 1.**
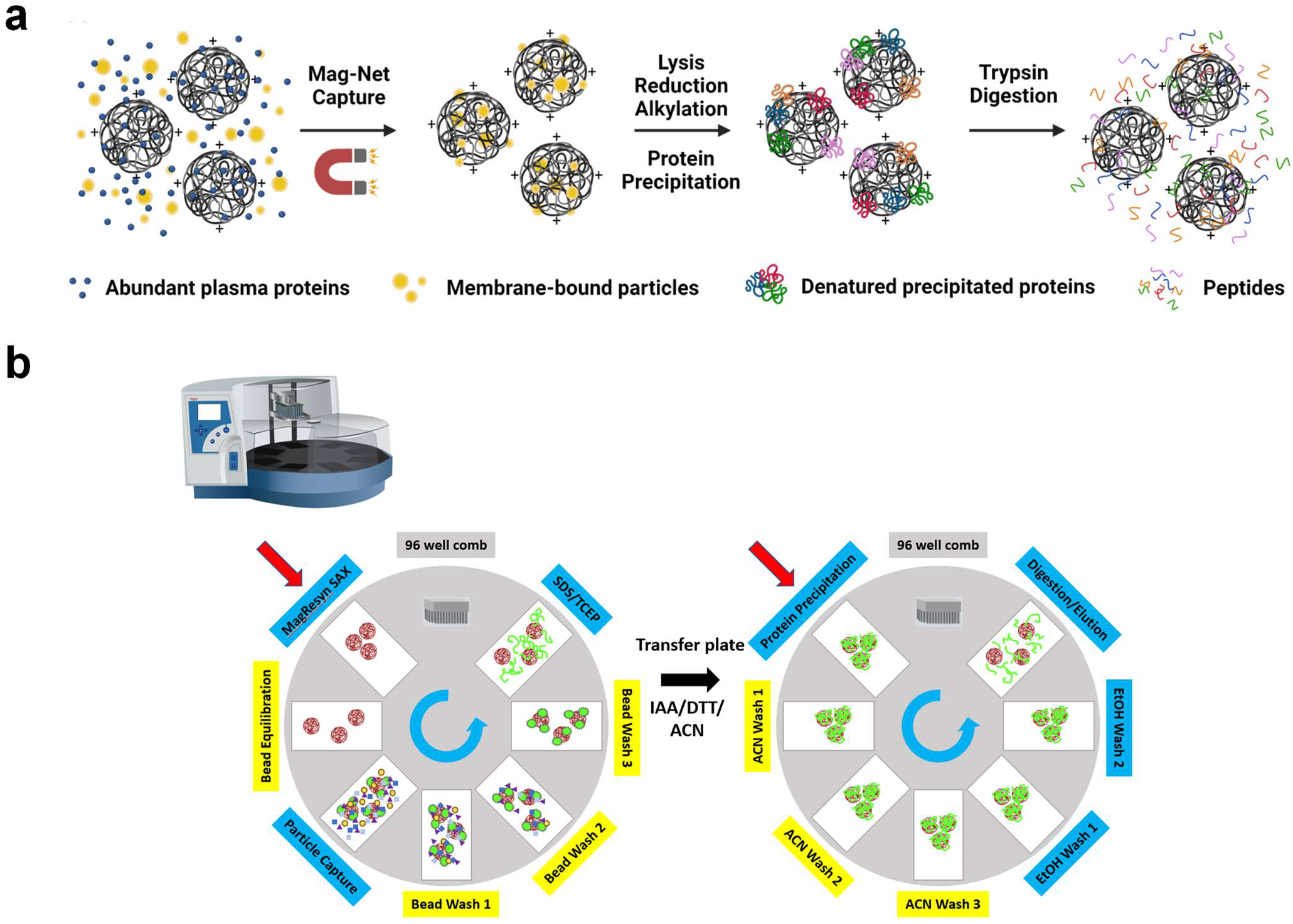
Mag-Net affinity capture workflow. a) SAX magnetic beads are incubated with plasma to sieve and bind negatively charged membrane-bound particles. The beads are immobilized with a magnet and abundant plasma proteins are gently washed away. The membrane particles are lysed on the beads and proteins are reduced and alkylated, precipitated back onto the beads, and then digested with trypsin. Peptides are collected and analyzed using nLC-MS/MS. b) The Kingfisher robot method to automate the Mag-Net workflow.

LC-MS/MS analysis of the Mag-Net enriched sample and the corresponding unfractionated plasma sample using data-independent acquisition mass spectrometry (DIA-MS) resulted in unusually large numbers of peptides and proteins that were detectable regardless of the computational method used to analyze the data (Figure 2). Using EncyclopeDIA v2.12.30 with a chromatogram library generated from six gas phase fractionated LC-MS/MS runs^30,31^ from the Mag-Net enriched sample, we were able to detect 4,163 protein groups in the enriched fraction and 1,088 protein groups from the corresponding unfractionated total plasma, prepared using standard PAC (Figure 2B, yellow bars). To demonstrate that the protein and peptide detections were not solely due to the use of an on-column chromatogram library generated using the Mag-Net enriched fraction, we also analyzed the data using EncyclopeDIA with a Prosit library^32^ generated from a human fasta file downloaded (September 2022) from Uniprot with one protein per gene (Figure 2B, blue bars).

**Figure 2.**
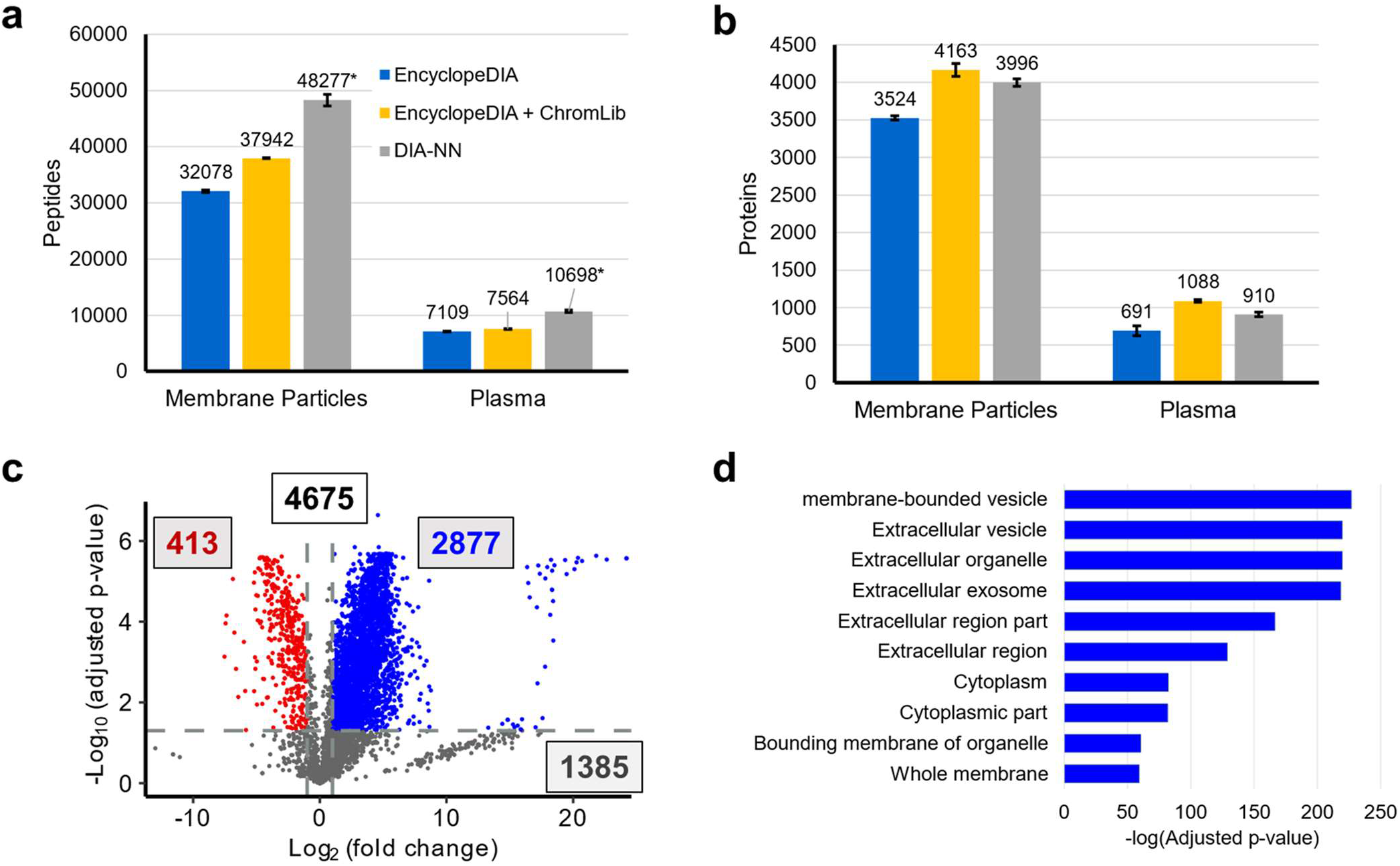
DIA-MS/MS analysis of the peptides from enriched membrane particles and unfractionated plasma. a) The number of peptides detected from three preparations of the same plasma sample at a 1% FDR with three different analysis strategies. The different analysis strategies were the use of EncyclopeDIA with a Prosit library (blue), EncyclopeDIA with a chromatogram library generated from gas phase fractionation (yellow), and DIA-NN in library free mode. EncyclopeDIA reports peptides detected and DIA-NN reports peptide precursors. The Mag-Net protocol detected a minimum of 451% more peptides than using the same analysis with unfractionated plasma. b) Proteins detected at a 1% FDR with the three different analysis strategies. The use of EncyclopeDIA with an on-column chromatogram library returned the most peptides and proteins and, thus, was used for all further analysis. c) Differential analysis of proteins between the enriched membrane particles (blue) and the same unfractionated plasma sample (red). d) The 2877 proteins that were enriched in the Mag-Net protocol were used to assess the enrichment of Jensen COMPARTMENTS using the Enrichr tool. * DIA-NN reports peptide precursors whereas EncyclopeDIA reports peptides. This makes the comparison of peptide detections between tools challenging.

Additionally, the analysis was also performed using DIA-NN in library free mode^33^ with the identical fasta file (Figure 2B, gray bars). The number of peptides (DIA-NN reports peptide precursors instead of peptides) detected in triplicate preparations and analysis of the membrane particle enriched fraction and unfractionated plasma are shown in Figure 2A. In all cases, the use of the MagReSyn® SAX beads to capture membrane particles prior to digestion resulted in improved coverage of the plasma proteome using a single LC-MS/MS analysis (Figure 2B). Additionally, the use of a chromatogram library generated from the gas phase fractionated analysis of the Mag-Net enriched sample improved the protein coverage of unfractionated plasma as described elsewhere^34^.

### Characterization of the proteins and particles enriched by Mag-Net

We performed a differential analysis of the proteins enriched in the membrane particle fraction versus the matched unfractionated plasma sample. In the analysis of the same sample prepared and measured six times, three times as unfractionated plasma and three times using the Mag-Net protocol, 4675 protein groups were detected. Of these, 2,877 proteins were enriched (BH-corrected p-value ≤ 0.05 and >1.5 fold change) in the membrane particle fraction, and 413 proteins were depleted from the unfractionated plasma (Figure 2C, Supplementary Figure S1). Statistical enrichment analysis using the Enrichr tool^35,36^ against the Jensen COMPARTMENTS resource^37^ revealed that the majority of the enriched proteins were proteins classically localized to membrane-bound vesicles, extracellular vesicles, exosomes, and extracellular organelles (Figure 2D)

To confirm that the fraction captured by Mag-Net is enriched in EVs, we calculated the fold enrichment of known extracellular vesicle proteins relative to the matched unfractionated plasma sample (Figure 3A). We used the known exosome marker proteins CD9, CD63, PDCD6IP (aka Alix), FLOT1, FLOT2, TSG101, SDCBP (aka Syntenin-1) and NCAM1^38–42^. NCAM has been used in the past for the immuno-enrichment and detection of neuronal derived exosomes from plasma or serum^43–46^. We used the proteins CD40, SEPTIN2, and ATP5F1A as markers of microvesicles^39,40,42,47,48^. The proteins HSP90B1 (aka Endoplasmin, GRP-94, or tumor rejection antigen 1) and ANXA5 (aka Annexin A5) were used as markers of cellular debris or apoptotic bodies^38,39,42,49,50^. Most of these markers were enriched >16 fold over their abundance in plasma and the canonical exosome markers CD9, CD63, Syntenin-1, and NCAM1 were all enriched >30-fold.

**Figure 3.**
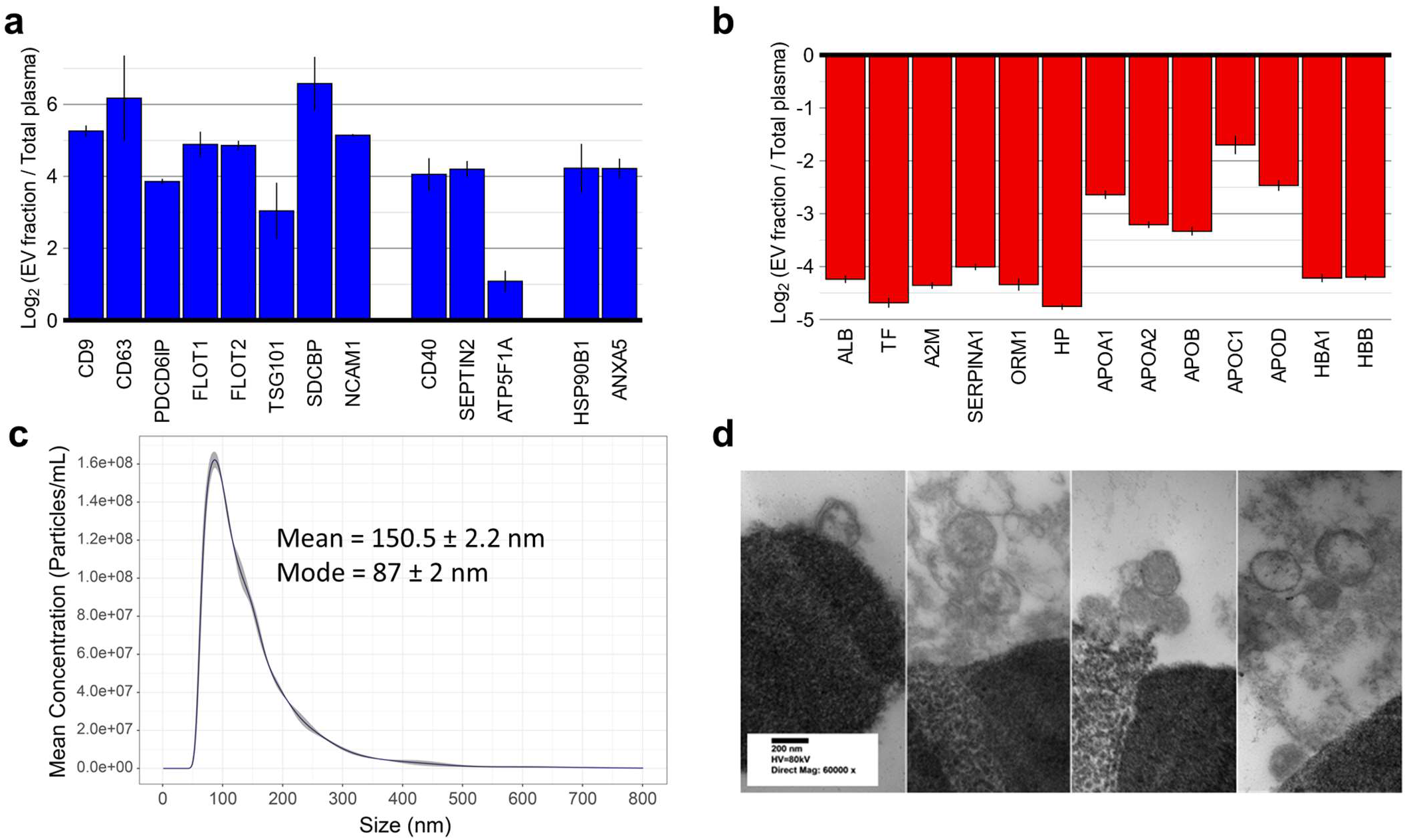
Characterization of Mag-Net enriched membrane particles. a) Fold enrichment of protein groups compared to total plasma. b) Fold depletion of protein groups compared to total plasma c) Nanosight particle count d) micrographs of captured membrane particles on bead surfaces using transmission electron microscopy.

This enrichment is a significant underestimate as many of these proteins are not detectable in plasma without enrichment and, thus, the quantity was estimated by integrating the background in the plasma sample where the peptide signal should be based on the Mag-Net sample. For example, the exosome marker CD9 had peptides that were enriched from 14.3 to 608.6-fold. The CD9 peptide with the lowest enrichment (DEPQRETLK) had a large background from a chemical interference in plasma, thus, an underestimate of the enrichment. We also confirmed the depletion of classic plasma and lipoprotein particle (LPP) markers from the Mag-Net fraction. The proteins albumin (ALB), serrotransferrin (TF), alpha-2-macroglobulin (A2M), alpha-1-acid glycoprotein 1 (ORM1), haptoglobin (HP), and hemoglobin (HBA1 and HBB) were all depleted by ∼95% -- a degree of depletion that is equal to or exceeds antibody depletion columns^6^.

A complication in the enrichment of plasma EVs is that lipoprotein particles are frequently co-enriched with EV fractions^19^. Lipoprotein particles are of similar size and density as extracellular vesicles, so EV enrichment methods using these physicochemical properties are particularly susceptible to contamination from lipoprotein particles. Mag-Net benefits from the MagReSyn® bead technology that facilitates the penetration of membrane particles and biomolecules throughout the volume of the polymer matrix. Thus, Mag-Net uses a combination of a size-based “netting” and charge-based binding from the quaternary ammonium surface chemistry. Using Mag-Net, lipoproteins were depleted by ∼80%, as determined by fold depletion of apolipoprotein protein groups (Figure 3b). Interestingly, APOB, the apolipoprotein present in low density lipoprotein (LDL) and very low density lipoprotein (VLDL) particles that has the closest diameter to extracellular vesicles, was depleted the most. The surface charge of EVs, where the surface is mostly phospholipid, has been shown to be significantly different from that of lipoprotein particles^51^. At the binding pH of 6.3, as performed in our analysis, the surface of lipoprotein particles will be significantly less negative than the EVs^51^ and therefore have significantly less affinity for the quaternary ammonium functionalized beads.

To verify that Mag-Net binds intact membrane particles we performed analyses to assess the particle size and morphology. After capture of particles to the MagReSyn® SAX bead we released the particles using salt and analyzed the fraction in triplicate using Nanoparticle Tracking Analysis (NTA) (Figure 3C). The most frequent particle size was 87 nm, with a mean size of 150 nm. Based on the distribution of the measured particle sizes, most were of a size that is typical of exosomes. However, NTA also confirmed that there are particles of size typically associated with microvesicles – confirming the results from the extracellular vesicle marker protein analysis (Figure 3A). Transmission electron microscopy (TEM) was also performed on the MagReSyn® beads and their captured membrane particles (Figure 3D). The ultrastructure images show vesicles with lipid bilayers captured on the surfaces of the MagReSyn® beads.

The Mag-Net protocol is unusual in that it combines the vesicle capture step and protein aggregation capture digestion step using a single magnetic bead (Figure 1). To assess whether the improved plasma protein coverage was because SAX beads were used in place of hydroxyl beads during the PAC digestion, we performed identical PAC digestion protocols with both types of beads. Supplementary Figure S2 shows a volcano plot of the protein differential abundance of the same plasma sample digested with the same protocol using either hydroxyl versus SAX functionalized magnetic beads. There are very few proteins with differential abundance between the two beads, confirming that the improved recovery was not because of the beads used during the PAC step. Importantly, these data also show that the SAX beads do not perform worse than the hydroxyl beads for PAC digestion.

To evaluate the “netting” capture ability of the MagReSyn® bead chemistry the Mag-Net vesicle capture protocol was also performed with the hydroxyl beads. The hydroxyl bead chemistry was chosen because those beads have an identical hyper-porous polymer matrix chemistry, but do not have the net cationic charge of the quaternary ammonium functionalized SAX beads. Supplementary Figure S3 is a volcano plot of the same plasma sample digested using PAC with hydroxyl beads versus performing the membrane capture protocol using the hydroxyl beads. Surprisingly, the uncharged hydroxyl functionalized beads also enriched proteins known to be present in membrane-bound vesicles and depleted abundant plasma proteins. To further understand the effect, we performed a differential analysis between the proteins enriched with the Mag-Net protocol using hydroxyl versus SAX beads (Supplementary Figure S4). In total, 3,334 protein groups were differentially abundant between the two bead types. While it was clear that the hydroxyl bead enriches for membrane bound particles when compared with an unenriched plasma digest, there were significantly more proteins enriched with the Mag-Net protocol performed using the SAX versus the hydroxyl beads – 2,992 and 342 proteins respectively (Supplementary Figure S4). Using the GO Cellular Component enrichment terms^52^ of the proteins enriched by the respective bead chemistry, the SAX beads enriched proteins (2,992) were derived from intracellular membranes, secretory granules, endosome membranes, etc. In contrast, the 342 proteins specific to the hydroxyl beads were from the lipoprotein particles, chylomicron particles, extracellular matrix, etc. This further confirms the importance of charge in the Mag-Net protocol in that it enables the enrichment of membrane vesicles while minimizing the contamination from much more abundant lipoprotein particles.

The challenges of the dynamic range of plasma have been described in detail previously^5,7^. We performed an analysis to evaluate the effect of Mag-Net on the dynamic range of measurements in plasma. The signal of each peptide was measured using the average of three replicate measurements performed using PAC total plasma digestion (Figure 4A) and the Mag-Net extracellular vesicle capture and digestionmethod (Figure 4B). If a peptide was not detected in a specific LC-MS run, the data from the runs where it was detected was used to provide boundaries to compute a background subtracted area where the peptide would be located if it were present. The peptide areas were normalized by equalizing the total ion current (TIC) between runs and the protein abundance calculated by summing the peptides mapping to the protein sequence. Figure 4 illustrates the significant enrichment of the EV marker proteins (blue) and the depletion of the abundant plasma proteins (red). The signal plotted for the protein abundance in the total plasma sample spans ∼10 logs (Figure 4A). It is important to point out that measuring the abundance for many of the proteins in the total plasma sample is only possible by matching the peptide signal from the respective measurement using the Mag-Net protocol. Despite the classic plasma proteins being depleted by ∼95% following Mag-Net (Figure 4B), those proteins are still some of the most abundant proteins in sample. This observation is analogous to what has been reported with the use of plasma depletion columns^10^. However, a notable difference is that the Mag-Net protocol depletes all proteins not associated with or within a membrane particle. Overall, Mag-Net significantly compresses the dynamic range of proteins in plasma without eliminating proteins.

**Figure 4.**
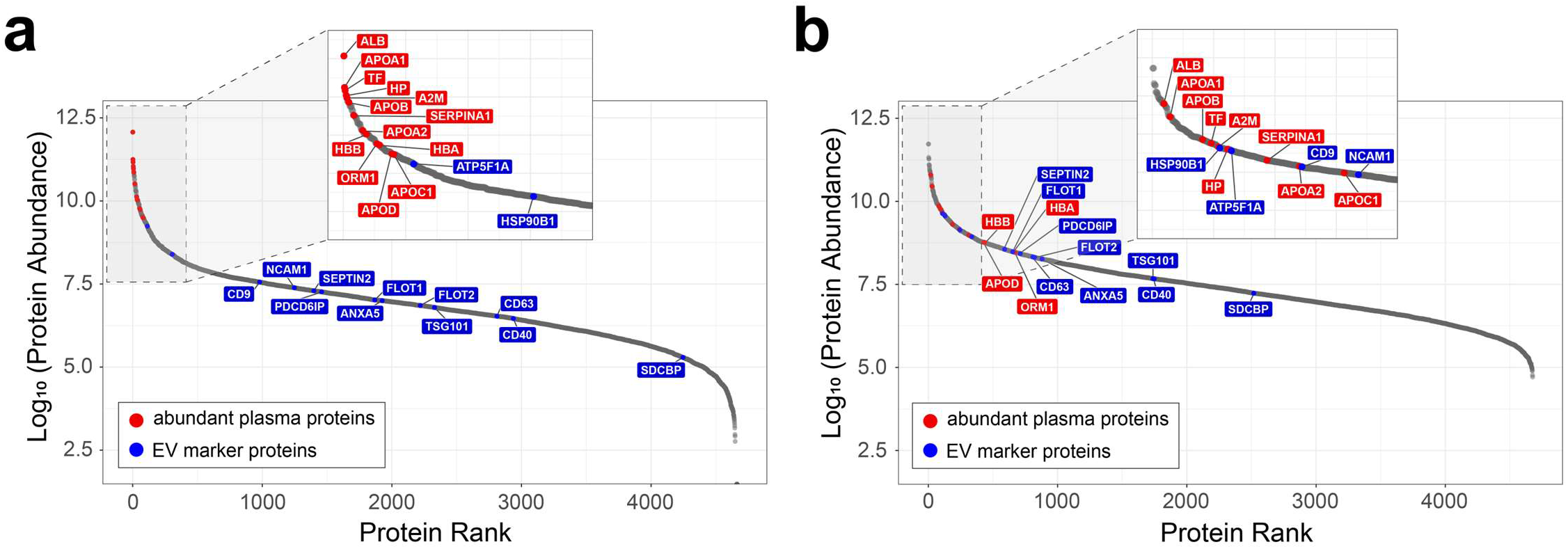
Dynamic range of the plasma proteome. The peptide signal was measured using the average of triplicate measurements performed using PAC total plasma digestion (A) and the Mag-Net extracellular vesicle capture method (B). If a peptide was not detected in a specific LC-MS run, the data from the runs where it was detected was used to provide boundaries to compute a background subtracted area where the peptide would be located if it were present. The peptide areas were normalized by equalizing the total ion current (TIC) between runs and the protein abundance calculated by summing the peptides mapping to a parsimonious protein group. The EV marker proteins are shown in blue and the classic abundant plasma proteins in red. The signal plotted for the protein abundance in the total plasma sample spans ∼10 logs. Despite the classic plasma proteins being depleted by ∼95% following Mag-Net, those proteins are still some of the most abundant proteins in sample.

### Assessment of quantitative accuracy and precision of samples prepared by Mag-Net

To confirm that the peptide measurements following Mag-Net enrichment reflected accurate quantities, we generated and analyzed matrix-matched calibration curves^53,54^. Three separate and independent matrix-matched calibration curves were made from the same human plasma sample diluted with chicken plasma. The calibration curve was generated by mixing human plasma with chicken plasma at 14 different volumetric ratios. The three separate matrix-matched calibration curves (14 dilution levels x 3 = 42 samples) were processed using the Mag-Net protocol. Each calibration curve was acquired using our standard DIA LC-MS protocol with runs ordered from lowest to highest proportion of human plasma. Several system suitability samples were performed as blanks between each calibration curve. LC-MS data were subjected to our normal analysis pipeline. The peptides that were conserved between human and chicken were removed and the peptide background-subtracted peak areas determined using Skyline^55^.

Figure 5 illustrates the analysis of a subset of the matrix-matched calibration curve^53^. The area of each peptide was divided by the area of the same peptide from the 100% undiluted human plasma sample. The measured log_2_ area ratios were plotted versus the expected % human plasma (Figure 5B). The distribution of the measured ratios is shown as both a box plot and a density plot. The mode of measured peptide relative abundance reflects the expected value based on the volumetric dilution prior to enrichment, sample preparation, and measurement. The median peptide ratios of the 1% human sample does underestimate the expected dilution. This indicates that a subset of the peptides is below the lower limit of quantification (LLOQ) at that dilution level.

**Figure 5.**
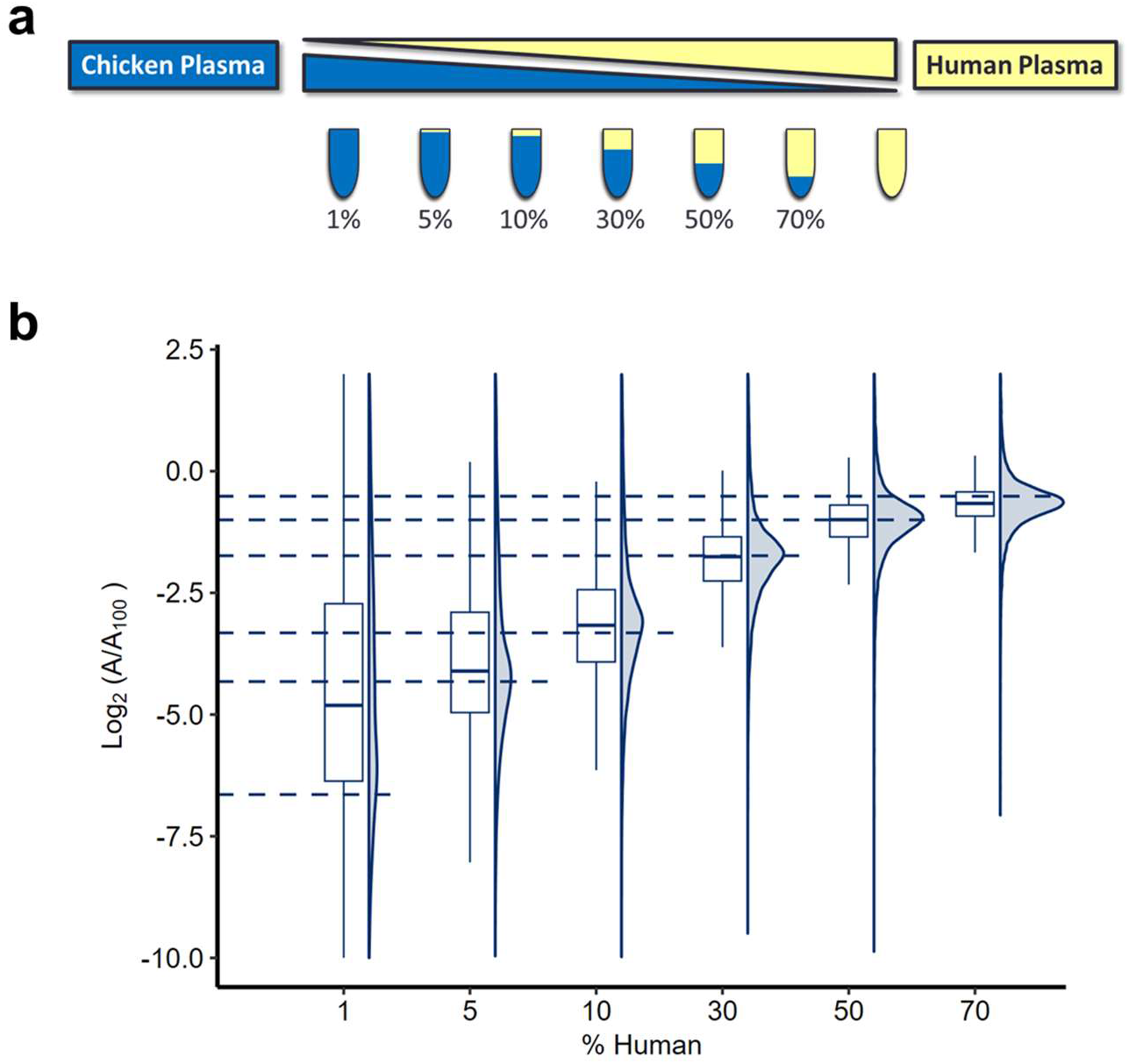
Summary of the matrix matched calibration curve data. A) Schematic of the volumetric mixing of human plasma with chicken plasma for 7 of the dilution points. The percentage represents the percent volume of human plasma where the remainder represents chicken plasma. The mixed plasma samples were enriched and digested using the Mag-Net protocol. Following peak detection with EncyclopeDIA, peptides conserved between human and chicken were removed. The peptide background subtracted peak areas were determined using Skyline and the ratio of the peptide area relative to the area from the 100% undiluted human sample. B) The measured Log_2_ area ratios versus the expected % human plasma.

We can also use these matrix-matched calibration curve data to calculate the LLOQ and the limit of detection (LOD) for each individual protein group and peptide. Supplementary Figure S5 contains representative plots of the calibration curves for two common EV markers (CD9 and NCAM1) and an abundant LDL protein marker (APOB). All three of these proteins have excellent quantitative linearity and an impressive LLOQ of 1.3%, 3.3% and 2.0% human plasma for CD9, NCAM1, and APOB respectively. Interestingly, even APOB, which is not enriched, but depleted by Mag-Net shows promising quantitative figures of merit. While every protein and peptide are different, a Skyline document containing the complete matrix matched calibration curve data is available at Panorama Public.

### Use of plasma EVs to assess molecular markers for Alzheimer’s Disease dementia (ADD)

EVs are membrane-bound structures that transport cargo between cells in the body. These important particles have diagnostic potential in several neurodegenerative diseases, including but not limited to ADD, PD, and amyotrophic lateral sclerosis. For example, proteins associated with exosomes (e.g. Alix, flotillins) are enriched in amyloid plaques in brain of people with of ADD^56^. The vesicle associated proteins Alix and Syntenin-1 are also important in the packaging of amyloid precursor protein and its amyloidogenic cleaved products into EVs^57^. EVs are involved in the extracellular enzymatic degradation of amyloid-β and promote both amyloid-β aggregation and clearance by microglia^58^. Synaptic proteins such as NPTX2, AMPA4, NLGN1, and NRXN2α have also been reported to be decreased in neuron derived EVs from plasma of patients with AD^59^. Even proteins like TDP-43, have been shown to be elevated in plasma derived exosomes of people diagnosed with ADD ^60^. Furthermore, recently it has been proposed that alpha-synuclein in plasma derived EVs is a potential diagnostic biomarker for PD^61–63^. It is also been reported that exosomes are responsible for transporting pathogenic forms of Tau between cells and throughout the periphery^64,65^. These prior studies indicate that the protein cargo in plasma EVs maybe a promising source of molecular markers for the diagnosis and monitoring of ADD and other neurodegenerative diseases.

To assess the potential of the Mag-Net method in the diagnostic capabilities of ADD and related diseases we performed a pilot study with plasma from 40 carefully selected individuals (Figure 6a). Most ADD biomarker studies include individuals with ADD and those who are age matched but otherwise healthy and cognitively normal. However, a complication with individuals with cognitive impairment is that they suffer from nonspecific lifestyle changes that can confound the specificity of any molecular signatures discovered. For example, individuals with dementia have altered eating habits and a change in appetite^66^, they are more sedentary and less physically active^67^, and socially isolated^68^. Moreover, individuals with ADD are commonly treated with two classes of drugs that temporarily provide symptomatic cognitive improvements. To control these confounding variables, we included a second group of individuals with dementia, those with PDD, who share nonspecific lifestyle changes of dementia and may be treated with the same symptomatic treatments but are commonly treated with different classes of medications for their movement disorder. We also included a third group of individuals diagnosed with PD but who were cognitively normal (PDCN); this group also is commonly treated for their movement disorder. The fourth group was healthy cognitively normal (HCN) individuals without any known neurodegenerative disease. These four groups, ADD, PDD, PDCN, and HCN, enable the detection of protein markers that change selectively based on the diagnostic group. We did our best to age match the individuals between groups and to maintain an equal male/female balance.

**Figure 6.**
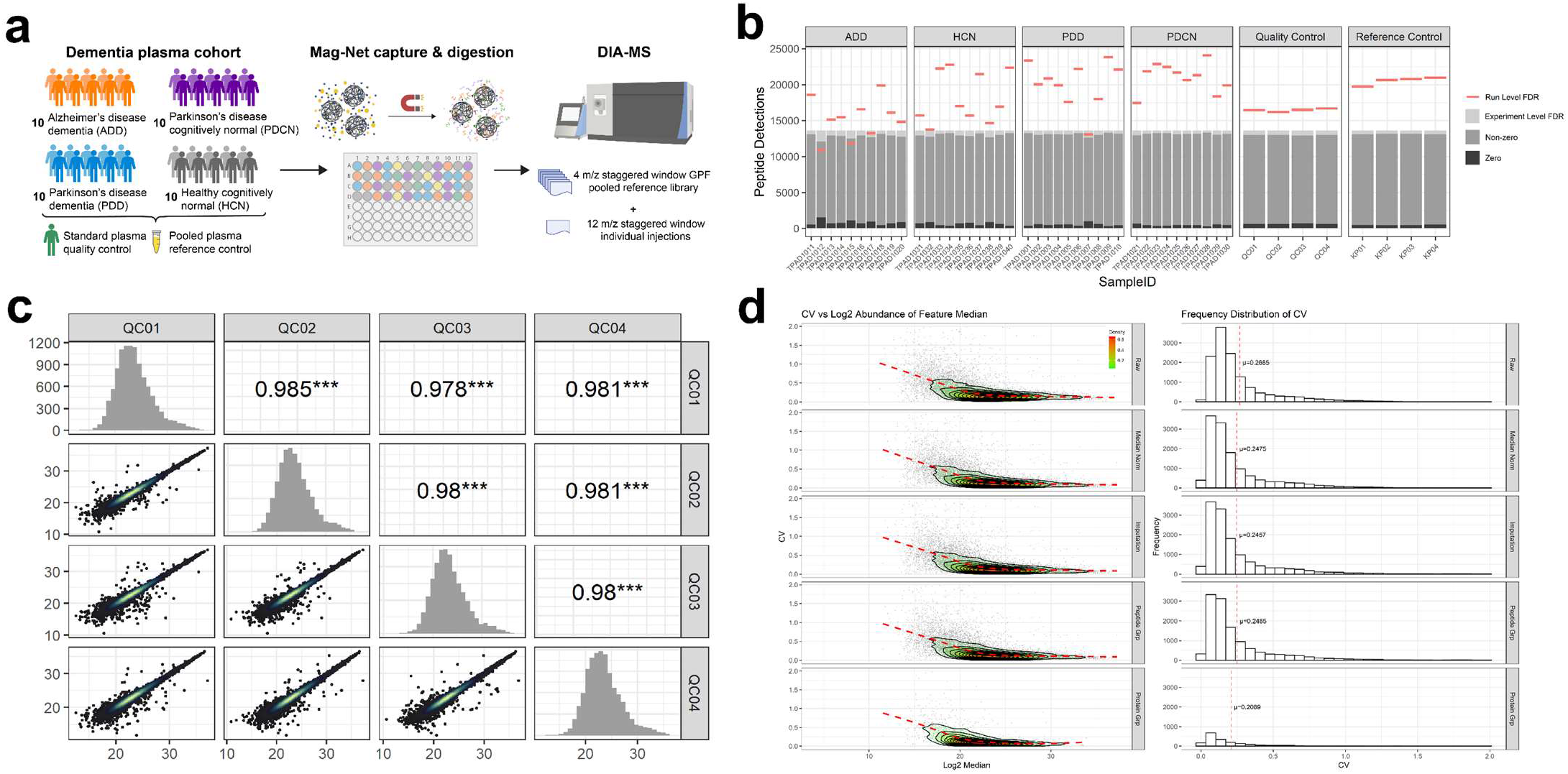
Application of Mag-Net for the assessment of markers of ADD, related diseases, and controls. a) We used a carefully selected cohort of plasma samples from 40 individuals (10 with ADD, 10 with PDD, 10 with PDCN, and 10 who were HCN). The individual samples in addition to two different controls were prepared using Mag-Net. The two pooled plasma sample were prepared four separated times and peptides quantified by data independent acquisition. One sample was used as a quality control for the quantitative measurements and the second was prepared as a reference control. b) The number of peptides detected when the FDR was controlled on the per run level (red lines) and on the entire experiment level (light grey). The dark grey represents the number of peptides with a non-zero quantity. c) The log2 peptide abundances measured between 4 separate preparations and measurement of the same quality control sample. The lowest Pearson correlation coefficient between pairs was 0.978. d) Effect of different data processing steps on the coefficient of variation of different peptide and protein quantities. The top row shows the RAW peptide intensities, followed by the data with median normalization, the effect of imputation, peptide grouping and protein grouping on the last row.

The measured peptide abundances are highly correlated between different preparations and analysis of the same sample. Figure 6c shows a plot of the unadjusted log2 peak area of the same peptide between runs. Replicate samples have Pearson correlation coefficients ranging from 0.978 to 0.985. The same data can also be assessed using a distribution of the coefficient of variation (CV) between the four replicates (Figure 6d). The mean (µ) CV of the RAW unadjusted areas between four preparations is 26.85%. After normalization, the median CV improved to 24.75%. Imputation of missing data values and peptide grouping did not significantly alter the CV. As highlighted previously^69^, protein grouping further improved the median CV to 20.89%

To assess whether individual protein quantities measured by Mag-Net could distinguish between disease states we used a combination of receiver operator characteristic (ROC) curve analysis and machine learning. We performed six pairwise analyses including 1) ADD versus all others, 2) HCN versus all others, 3) PDD versus all others, 4) PDCN versus all others, 5) dementia (ADD and PDD) versus cognitively normal (HCN and PDCN), and 6) Parkinson’s disease (PDCN and PDD) versus non-Parkinson’s disease (ADD and HCN). The total analysis is summarized in Supplementary Table 1. Using a two-sided Mann-Whitney U test we found 204 individual proteins that distinguished ADD from all others after multiple hypothesis correction (q-value < 0.05) and 310 proteins that distinguished patients with PD (PDD or PDCN) from those with ADD or HCN. Despite successfully finding individual proteins that were unique to ADD or PD, we found 0 proteins that were specific to PDD, PDCN, HCN, or to patients with dementia in general (both ADD and PDD). These findings were not surprising and highlight the limitations of a relatively small sample size. Additionally, despite finding proteins that can distinguish between PD and non-PD patients, we expected that PDD and PCN would be more challenging because they share underlying Lewy body pathology. We also recognize that our older HCN cohort, age matched to our ADD participants, might not be ideally matched to PD groups.

In total, there were 1093 proteins that had an ROC > 0.7 from the six pairwise analyses described above. These proteins are summarized in the heatmap in Figure 7a. Unsupervised clustering of these proteins shows that these proteins separate the patients with ADD from the PD patients (both PDD and PDCN).

**Figure 7.**
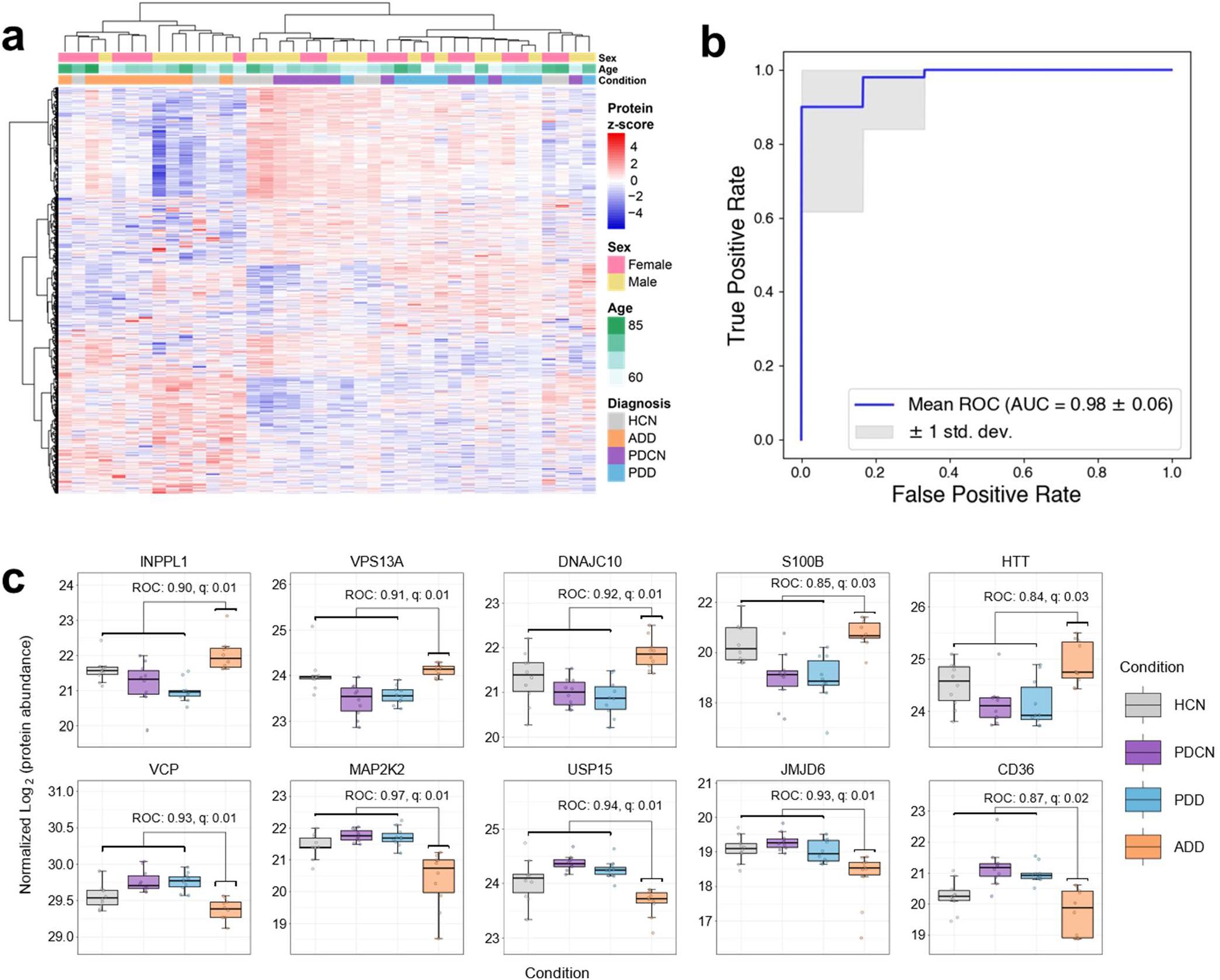
Measurement of proteins that provide separation between disease cohorts. a) Heatmap of 1093 proteins that had an ROC > 0.7 from any of the six pairwise analyses described in the text. b) A receiver operator characteristic (ROC) curve that illustrates our ability to correctly separate people with ADD from other groups (i.e. PDD, PDCN, and HCN) using a hard-margin SVM with linear kernel. The plot shows the mean and standard deviation of 10 iterations of five-fold stratified cross validation with different random splits. c) Illustration of selected protein coding genes that are either increased (INPPL1, VPS13A, DNAJC10, S100B, and HTT) or decreased (VCP, MAP2K2, USP15, JMJD6, and CD36) in ADD plasma when compared with the other three disease groups.

We manually examined proteins that distinguished ADD from the other three patient groups. Figure 7c highlights 10 proteins – five that are increased and five that are decreased in ADD, all of which have been implicated in the pathogenesis of AD. The protein encoded by the INPPL1 gene (aka SHIP2) is a phosphoinositide 5-phosphatase (PI 5-phosphatase) that is increased relative to the other conditions, and these key intracellular signaling molecules are implicated in the pathogenesis of AD^70^. In particular, the transcript of INPPL1 in the brain has been shown to correlate with cognitive decline of ADD patients^71^. The protein encoded by INPPL1 is a mediator of amyloid-beta toxicity by the regulation of tau hyperphosphorylation^72^ and actin cytoskeleton reorganization^73^. VPS13A is a protein that localizes to the membrane contact site between the endoplasmic reticulum (ER) and mitochondria. VPS13A is known to mediate the transfer of lipids and is required for efficient lysosomal degradation. Endolysosomal abnormalities, mitochondrial dysfunction, ER stress, and altered lipid metabolism are commonly observed in both AD and PD. The protein encoded by the gene DNAJC10 (aka DJC10) is an ER quality control protein, and its transcript has been shown to be both associated and predictive of ADD^74^. The protein S100B is part of RAGE signaling and activates small GTPases such as Ras and Rac1. S110B is known to be elevated in the plasma of ADD patients^75^ and has been implicated in the mechanisms underlying neurodegeneration in ADD^76^ and PD pathogenesis^77^. Serum levels of S100B have been proposed as a marker to distinguish the severity and progression of ADD^75^ as it could provide a measure of astrocytic reaction to neuronal injury. Another elevated protein in the ADD cohort is Huntingtin (aka HD) encoded by the gene HTT. HTT had an ROC = 0.84 (q-value 0.03). Huntingtin levels are known to be elevated in the hippocampus of ADD patients and has been shown to co-aggregate with amyloid-beta plaques^78^. There were several other proteins known to be involved in vesicle-mediated transport and lipid metabolism that were increased in ADD patients that are highlighted in Supplementary Figure S7 and S8.

Figure 7c illustrates five proteins that are decreased in abundance in ADD. The protein encoded by the gene VCP (Valosin-containing protein aka TERA) is an ER ATPase that is responsible for removing tau fibrils^79^. Mutations in VCP have been shown to be associated with late-onset ADD^80^. The protein colocalizes in the affected neurons of PD and amyotrophic lateral sclerosis (ALS)^81^. MAP2K2 is a MAP kinase (MAPK-activated protein kinase 2), activates ERK^82^, is involved in neuroinflammation and neurodegenerative pathology, and overexpressed in mouse APP and PSEN1 models^83^. USP15 mediates deubiquitination of substrates bound to VCP^84^ and antagonizes Parkin-mediated mitochondrial ubiquitination and mitophagy^85^. JMJD6 has been identified as one of the key potential neuronal drivers in AD^86^. The protein CD36 acts as a scavenger receptor for amyloid fibrils, and as the disease progresses, microglial cells are known activate into a pro-inflammatory phenotype and downregulate CD36 expression^87^. Additionally, we noticed many proteins known to be involved in ubiquitin proteasome mediated protein degradation were decreased (Supplementary Figure S9). Other proteins that decreased are involved in intracellular protein transport (e.g. proteins with GTPase activity) and in microtubule-based transport (e.g. kinesins, dynactin complex members, and cadherin binding).

While there are many proteins that show an altered abundance between ADD and the other three diagnosis groups, we wanted to evaluate our ability to create a machine learning classifier that could distinguish between groups. Six separate binary classifiers were trained on the normalized and batch corrected data to predict each of the four diagnostic groups individually (ADD, PDD, PDCN, HCN) as well as Parkinson’s disease (PDD and PDCN) and dementia (ADD and PDD). We use a hard-margin support vector machine (SVM) with linear kernel for classification because it is both inherently interpretable and well suited for small, high-dimensional datasets^88^. To estimate the generalization performance of the learned classifiers on each task, we performed 10 iterations of five-fold stratified cross validation with different random splits and report the mean and standard deviation of the area under the validation ROC curve across folds. Figure 7b is a receiver operator characteristic (ROC) curve that illustrates our ability to correctly separate people with ADD from other groups (i.e. PDD, PDCN, and HCN). The results of the other binary classifiers are illustrated in Supplementary Figure S10. All binary classifiers perform very well, except for the classifier attempting to identify HCN individuals from those with either ADD, PDD, or PDCN. The relatively poor performance of our HCN classifier could be explained by a percentage of the patients having undiagnosed cognitive impairment.

## CONCLUSION

The Mag-Net protocol is a simple yet powerful approach for the analysis of the plasma proteome. The enrichment of proteins within EVs enables the detection of proteins that would otherwise be beyond the dynamic range of LC-MS/MS in unfractionated plasma. Mag-Net benefits from the MagReSyn® bead technology that facilitates the penetration of membrane particles and biomolecules throughout the volume of the polymer surface. Thus, Mag-Net uses both a size based “netting” and charge-based binding from the quaternary ammonium surface chemistry. Mag-Net enables the selective enrichment of vesicles in the presence of similar size lipoprotein particles.

Combined with a KingFisher magnetic bead robot, Mag-Net enables the automated membrane particle capture, lysis, and subsequent aggregation of vesicle proteins on a single bead. Performing PAC on the same beads used for the vesicle enrichment results in excellent recovery and minimal losses since efficient elution of the vesicles from the beads is unnecessary. Instead, peptides are released from the beads via tryptic digestion. Because samples are prepared in a 96-well format, up to 96 samples can be prepared in parallel. The Mag-Net protocol is particularly well suited for high throughput applications requiring the preparation of many plasma samples for analysis by LC-MS/MS.

The Mag-Net protocol enables the reproducible and accurate measurement of >37,000 peptides and >4,000 proteins from human plasma when combined with data independent acquisition mass spectrometry. The correlation coefficients from pairwise comparisons of the same sample prepared and measured multiple times was >0.978. The median CV of peptide and protein measurements between preparations was 24.75% and 20.89% respectively. The peptide measurements showed excellent accuracy for many peptides up to 100:1 dilution using a matrix-matched calibration curve. This simple, robust, and cost-effective solution may enable the identification of novel markers of disease diagnosis, progression, and response to therapy.

We demonstrate that many of the pathways and functional categories of proteins known to be disrupted in ADD in the brain are also altered in plasma EVs. While it is unlikely that all these proteins in plasma are derived from the brain or central nervous system (CNS) it suggests that changes in this subset of the plasma proteome can provide a signature for biochemical and cellular changes that are occurring in neurodegenerative disease.

## METHODS

### Plasma samples

For method optimization and validation, plasma samples from 17 individual healthy donors collected in K_2_EDTA vials were purchased from Innovative Research (IPLASK2E2ML, Innovative Research Inc, Novi, MI). Samples were aliquoted and stored at −80°C. Plasma samples previously banked at Stanford as part of longitudinal studies on neurodegenerative diseases of aging were selected. Diagnostic criteria for ADD and PD and the assessments for the cognitive status determination have previously been described^89^. Ten age- and sex-matched samples were selected from each of four categories: 1) Alzheimer’s disease dementia (ADD), 2) Parkinson’s disease with dementia (PDD), 3) Parkinson’s disease cognitively normal (PDCN), and 4) healthy cognitively normal (HCN). Study protocols were approved by the institutional review board of Stanford University and written informed consent was obtained from all participants or their legally authorized representative.

### Membrane particle enrichment coupled with protein aggregation capture (Mag-Net Method)

Membrane particle capture from plasma and protein aggregation capture (PAC) steps were performed on a Thermo Scientific™ KingFisher Flex. The combination of the membrane particle capture with PAC and digestion is referred to as Mag-Net (Note: Eight plates were designated for each of two Kingfisher (KF) cycles as follows. KF Cycle 1: Plate 1: SAX beads, Plate 2: Bead Equilibration, Plate 3: Membrane Particle Capture, Plate 4-6: Wash, Plate 7: Solubilization/Reduction, Position 8: 96 well comb. KF Cycle 2: Plate 1: Same as Plate 7 of KF Cycle 1, Plate 2-4: Acetonitrile Wash, Plate 5-6: Ethanol Wash, Plate 7: Trypsin Digestion/Peptide Elution.) Briefly, HALT cocktail (protease and phosphatase inhibitors, Thermo Fisher Scientific) was added to plasma (20-100 µL) and then mixed 1:1 by volume with Binding Buffer (BB, 100 mM Bis-Tris Propane, pH 6.3, 150 mM NaCl). MagReSyn® strong anion exchange or MagResyn® Hydroxyl magnetic microparticles (ReSyn Biosciences) were first equilibrated 2 times in Equilibration/Wash Buffer (WB, 50 mM Bis Tris Propane, pH 6.5, 150 mM NaCl) with gentle agitation and then combined in a 1:4 ratio (volume beads to volume starting plasma) with the plasma:BB sample for 45 minutes at room temperature. The beads were washed with WB 3 times for 5 minutes with gentle agitation. The enriched membrane particles on the beads were then solubilized and reduced in 50 mM Tris, pH 8.5/1% SDS/10 mM Tris (2-carboxyethyl) phosphine (TCEP) with 800 ng enolase standard added as a process control.

Following reduction, the plate was removed from the Kingfisher. Samples were alkylated with 15 mM iodoaceamide in the dark for 30 minutes and then quenched with 10 mM DTT for 15 minutes. Total unfractionated plasma (1 µL) was prepared in parallel (reduction, alkylation, and quenched) and added to the Kingfisher plate as a control for enrichment. The samples were processed using protein aggregation capture with minor modifications. Briefly, the samples were adjusted to 70% acetonitrile, mixed, and then incubated for 10 minutes at room temperature to precipitate proteins onto the bead surface. The beads were washed 3 times in 95% acetonitrile and 2 times in 70% ethanol for 2.5 minutes each on magnet. Samples were digested for 1 hour at 47 °C in 100 mM ammonium bicarbonate with Thermo Scientific™ Pierce™ porcine trypsin at a ratio of 1:20 trypsin to protein. The digestion was quenched to 0.5% trifluoracetic acid and spiked with Pierce Retention Time Calibrant (PRTC) peptide cocktail (Thermo Fisher Scientific) to a final concentration of 50 fmol/µL. Peptide digests were then stored at −80°C until LC-MS/MS analysis.

### Matched matrix calibration sample preparation

Human plasma was diluted with chicken plasma by volumetric mixing for 14 human:chicken dilution points (100:0, 70:30, 50:50, 30:70, 10:90, 7:93, 5:95, 3:97, 1:99, 0.7:99.3, 0.5:99.5, 0.3:99.7, 0.1:99.9, 0:100). Membrane particles were enriched using the Mag-Net Method for each dilution and performed in triplicate.

### Nanoparticle tracking analysis

Membrane particles were eluted from SAX beads in Elution Buffer (25 mM Bis Tris Propane, pH 6.5, 1 M NaCl, 0.1% Tween 20) and diluted in PBS (first 1:50 and then an additional dilution of either 1:25 or 1:50). Diluted samples were analyzed using the Nanosight NS300 (Malvern Panalytical Ltd). The instrument was calibrated using a NIST standardized 100 nm latex bead and 5% CV rate. Using a manual injection method, videos were captured for each sample at 21.7°C. Particle counts were determined using the Nanosight Nanoparticle Tracking Analysis (NTA) software 2.3 (Malvern Panalytical Ltd) with a detection threshold of 3-Multi.

### Transmission electron microscopy (TEM)

The samples (membrane particles bound to SAX beads) were fixed in 2% glutaraldehyde/0.1 M sodium cacodylate, pH 7.2 overnight at 4°C, and then postfixed in 2% OsO_4_ buffered with 100 mM sodium cacodylate, pH 7.2 overnight at room temperature. The samples were then washed with water, dehydrated in an increasing ethanol series (30%, 50%, 60%, 70%, 80%, 90%, 98% for 10 minutes each followed by 100% x 3 for 10 minutes) and then infiltrated stepwise from ethanol into Spurr’s resin (2:1, 1:1, 1:2 for 30 min each followed by 100% Spurr’s x 3 overnight) and then embedded and baked at 60°C for 2 days. Ultra-thin sections were cut from each sample, collected on grids, and imaged on a JEOL 1230 Transmission Electron Microscope (Peabody, MA).

### Liquid chromatography and mass spectrometry

One µg of each sample with 150 femtomole of PRTC was loaded onto a 30 cm fused silica picofrit (New Objective) 75 µm column and 3.5 cm 150 µm fused silica Kasil1 (PQ Corporation) frit trap loaded with 3 µm Reprosil-Pur C18 (Dr. Maisch) reverse-phase resin analyzed with a Thermo Easy-nLC 1200. The PRTC mixture is used to assess system suitability before and during analysis. Four of these system suitability runs are analyzed prior to any sample analysis and then after every six sample runs another system suitability run is analyzed. Buffer A was 0.1% formic acid in water and buffer B was 0.1% formic acid in 80% acetonitrile. The 40-minute system suitability gradient consists of a 0 to 16% B in 5 minutes, 16 to 35% B in 20 minutes, 35 to 75% B in 1 minute, 75 to 100% B in 5 minutes, followed by a wash of 9 minutes and a 30-minute column equilibration. The 110-minute sample LC gradient consists of a 2 to 7% for 1 minute, 7 to 14% B in 35 minutes, 14 to 40% B in 55 minutes, 40 to 60% B in 5 minutes, 60 to 98% B in 5 minutes, followed by a 9-minute wash and a 30-minute column equilibration. Peptides were eluted from the column with a 40°C heated source (CorSolutions) and electrosprayed into a Thermo Orbitrap Eclipse Mass Spectrometer with the application of a distal 2 kV spray voltage.

For each batch of samples, we collected a chromatogram library using gas phase fractionation from 6 independent injections using a pool of all samples within the batch. These gas phase fractionation runs collected tandem mass spectrometry data each covering a different portion of the total m/z range. For each injection MS1 spectra were collected with a mass range of 100 *m/z* (400-500 *m/z*, 500-600 *m/z*…900-1000 *m/z*). MS2 spectra were collected using the tMSn scan function with a loop count of 25 at 30,000 resolving power, a normalized AGC target of 1000%, maximum injection time set to Auto, and 27% normalized collision energy. Each MS2 spectrum was collected using a 4 m/z isolation window with the edges placed in peptide forbidden zones as described previously^30,31^. In alternating cycles of data independent acquisition, the location of the isolation windows were offset by 50% so that the data could be computationally demultiplexed to 2 m/z^90^. The chromatogram library data is used to quantify proteins from individual sample runs.

Each sample for quantitative analysis was run once using consist of a cycle of one 30,000 resolving power full-scan mass spectrum with a mass range of 395-1005 m/z, AGC target of 4e5, maximum injection time set to Auto. MS2 spectra were collected using the tMSn scan function, loop count of 75 at 30,000 resolving power, normalized AGC target of 800%, maximum injection time set to Auto (54 ms), and 27% normalized collision energy. Each MS2 spectrum was collected using staggered 12 m/z isolation windows with optimized window placements^31^. Application of the mass spectrometer and LC solvent gradients are controlled by the ThermoFisher Xcalibur

### Peptide detection and quantitative signal processing

Thermo RAW files were converted to mzML format using Proteowizard (version 3.0.23063) using vendor peak picking and demultiplexing with the settings of “overlap_only” and Mass Error = 10.0 ppm^90^. On column chromatogram libraries were created using the data from the six gas phase fractionated “narrow window” DIA runs of the pooled reference as described previously^30,31^. These narrow windows were analyzed using EncyclopeDIA (v2.12.30 with version 2 scoring) with the default settings (10 ppm tolerances, trypsin digestion, HCD b- and y-ions) of a Prosit predicted spectra library based the Uniprot human canonical FASTA (September 2022). The results from this analysis from each batch were saved as a “Chromatogram Library’’ in EncyclopeDIA’s eLib format where the predicted intensities and iRT of the Prosit library were replaced with the empirically measured intensities and RT from the gas phase fractionated LC-MS/MS data. The “wide window” DIA runs were analyzed using EncyclopeDIA (v2.12.30 with version 2 scoring) requiring a minimum of 4 co-varying and interference free quantitative transitions and filtering peptides with q-value ≤ 0.01 using Percolator 3.01. After analyzing each file individually, EncyclopeDIA was used to generate a “Quant Report’’ which stores all detected peptides, integration boundaries, quantitative transitions, and statistical metrics from all runs in an eLib format. The Quant Report eLib library is imported into Skyline-daily with the human uniprot FASTA as the background proteome to map peptides to proteins, perform peak integration, manual evaluation, and report generation. A csv file of peptide level total area fragments (TAFs) for each replicate was exported from Skyline using the custom reporting capabilities of the document grid. To examine enrichment and depletion relative to total, the group comparison document grid was used to export a csv file of protein-level average ratios, maximum/minimum ratios, and p-values. Protein-level matrix-matched calibration curve data was exported as a csv file of protein abundance for each replicate.

Despite precautions taken to ensure equivalent sample preparation, handling and acquisition, additional post-processing was performed to normalize, and batch adjust the quantitative data to remove residual technical noise. We performed protein parsimony as described previously^91^ and summed the peptide for each protein group. Modeling the proportional changes of protein group intensities, log2 transformation is applied followed by equalizing the medians between to the protein group values across instrument runs under the assumption that median total area fragment values should be equal sans batch effect from known and unknown sources of variability. We used the limma “removeBatchEffect” function to minimize batch effects derived from run order and also the effect of “cohort”.

### Development of a classifier to differentiate between ADD, PDD, PDCN, and HCN individuals

Six separate binary classifiers were trained on the normalized and batch corrected data to predict each of the four diagnostic groups individually (ADD, PDD, PDCN, HCN) as well as Parkinson’s disease (PDD and PDCN) and dementia (ADD and PDD). We use a hard-margin support vector machine (SVM) with linear kernel for classification because it is both inherently interpretable and has properties well suited for small, high-dimensional datasets^88^. To estimate the generalization performance of the learned classifiers on each task, we performed 10 iterations of 5-fold stratified cross validation with different random splits and report the mean and standard deviation of the validation ROC curve across folds.

In addition to our SVM classifiers, we also evaluated the predictive power of each of the 2343 measured proteins individually. The area under the ROC curve was calculated for each protein on each of the six prediction tasks, and significance assigned using a two-sided Mann-Whitney U test. The resulting p-values were corrected using the Benjamini-Hochberg procedure for controlling the false discovery rate^92^, yielding a q-value for each putative biomarker.

## DATA AVAILABILITY

All mass spectrometry RAW datafiles, demultiplexed mzML intermediate files, EncyclopeDIA output, and final Skyline documents are available in Panorama Public at https://panoramaweb.org/Mag-Net.url. The dataset is registered through the ProteomeXchange with the unique identifier (PENDING). R scripts used to produce figures in the manuscript and the methods for the KingFisher robot are available at https://github.com/uw-maccosslab/Mag-Net

## ACKNOWLEDGEMENTS

We would like to thank members of the Hoofnagle and MacCoss labs for critical discussions. We would like to thank the Vision Core Lab for providing access to the JEOL 1230 TEM. This work is supported in part by the National Institutes of Health grants U19 AG065156, P30 AG013280, U01 DK137097, R01 NS115114, P30 AG066515, F31 AG069420, and T32 AG066574.

## AUTHOR CONTRIBUTIONS

CCW and MJM conceived the study and designed the method validation experiments. KP, TJM, CCW, and MJM designed the pilot experiment and controls. KP and TJM selected plasma specimens from retrospective ADD and PD collections and aided in the data interpretation. CCW developed and optimized the Mag-Net method. PN, SB, IG, SS and JJ provided expertise, manufactured non-commercial MagReSyn® bead formulations, contributed to scientific interpretation, and assisted with inter-laboratory method evaluation. KT, JP, DP, GM, and MR contributed to data analysis and generated data figures. AH contributed scientific interpretation. EH and EP provided technical support. CCW and MJM wrote the manuscript. All authors read and provided comments about the manuscript.

## COMPETING FINANCIAL INTERESTS

Ireshyn Govender, Stoyan Stoychev and Justin Jordaan are employed by ReSyn Biosciences, proprietors of MagReSyn® technology. The MacCoss Lab at the University of Washington has a sponsored research agreement with Thermo Fisher Scientific, the manufacturer of the mass spectrometry instrumentation used in this research. Additionally, Michael MacCoss is a paid consultant for Thermo Fisher Scientific.

## SUPPLEMENTARY DATA (Mag-Net EV Method Supplement.pdf)

- Figure S1: Volcano plot of human plasma versus Mag-Net enriched EVs with representative markers highlighted.
- Figure S2: Volcano plot comparing the use of MagReSyn Hydroxyl versus SAX beads for protein aggregation capture (PAC) digestion.
- Figure S3: Volcano plot comparing the preparation of total plasma using PAC with Hydroxyl beads versus the use of Mag-Net membrane particle capture with Hydroxyl beads.
- Figure S4: Volcano plot comparing the difference between the use of MagResyn Hydroxyl versus SAX beads for Mag-Net.
- Figure S5: Example matrix matched calibration curves for the EV marker proteins CD9 and NCAM1 and the (V)LDL marker APOB.
- Figure S6: Assessment of plasma freeze thaw cycles on the relative peptide abundance.
- Figure S7: Selected proteins elevated in ADD that are known to be involved in vesicle mediated transport.
- Figure S8: Selected proteins elevated in ADD that are known to be involved in lipid metabolism.
- Figure S9: Selected proteins decreased in ADD that are known to be involved in ubiquitin proteosome mediated protein degradation.
- Figure S10: Individual ROC curve analysis illustrating the performance of a trained hard-margin support vector machine (SVM) with a linear kernel.

## Notes

https://panoramaweb.org/Mag-Net.url

https://github.com/uw-maccosslab/Mag-Net

### Summary of Updates

Added an analysis of a small cohort of plasma samples from patients with neurodegenerative disease.

## REFERENCES

1. Bruderer, R. et al. Analysis of 1508 Plasma Samples by Capillary-Flow Data-Independent Acquisition Profiles Proteomics of Weight Loss and Maintenance ’[S]. Molecular & Cellular Proteomics 18, 1242–1254 (2019).

2. Liu, Y. et al. Quantitative variability of 342 plasma proteins in a human twin population. Mol. Syst. Biol. 11, 786 (2015).

3. Geyer, P. E., Holdt, L. M., Teupser, D. & Mann, M. Revisiting biomarker discovery by plasma proteomics. Molecular Systems Biology 13, 942 (2017).

4. Niu, L. et al. Plasma proteome profiling discovers novel proteins associated with non-alcoholic fatty liver disease. Mol Syst Biol 15, e8793 (2019).

5. Anderson, N. L. & Anderson, N. G. The Human Plasma Proteome: History, Character, and Diagnostic Prospects*. Molecular & Cellular Proteomics 1, 845–867 (2002).

6. Shi, T. et al. IgY14 and SuperMix immunoaffinity separations coupled with liquid chromatography– mass spectrometry for human plasma proteomics biomarker discovery. Methods 56, 246–253 (2012).

7. Anderson, N. L. The clinical plasma proteome: a survey of clinical assays for proteins in plasma and serum. Clin Chem 56, (2010).

8. Pieper, R. et al. Multi-component immunoaffinity subtraction chromatography: an innovative step towards a comprehensive survey of the human plasma proteome. Proteomics 3, 422–432 (2003).

9. Keshishian, H., Addona, T., Burgess, M., Kuhn, E. & Carr, S. A. Quantitative, Multiplexed Assays for Low Abundance Proteins in Plasma by Targeted Mass Spectrometry and Stable Isotope Dilution*. Molecular & Cellular Proteomics 6, 2212–2229 (2007).

10. Tu, C. et al. Depletion of abundant plasma proteins and limitations of plasma proteomics. J Proteome Res 9, 4982–4991 (2010).

11. Keshishian, H. et al. Multiplexed, Quantitative Workflow for Sensitive Biomarker Discovery in Plasma Yields Novel Candidates for Early Myocardial Injury*. Molecular & Cellular Proteomics 14, 2375–2393 (2015).

12. Addona, T. A. et al. A pipeline that integrates the discovery and verification of plasma protein biomarkers reveals candidate markers for cardiovascular disease. Nat Biotechnol 29, 635–643 (2011).

13. Qian, W.-J. et al. Enhanced Detection of Low Abundance Human Plasma Proteins Using a Tandem IgY12-SuperMix Immunoaffinity Separation Strategy. Mol Cell Proteomics 7, 1963–1973 (2008).

14. Tognetti, M. et al. Biomarker Candidates for Tumors Identified from Deep-Profiled Plasma Stem Predominantly from the Low Abundant Area. J. Proteome Res. 21, 1718–1735 (2022).

15. Blume, J. E. et al. Rapid, deep and precise profiling of the plasma proteome with multi-nanoparticle protein corona. Nat Commun 11, 3662 (2020).

16. Ferdosi, S. et al. Engineered nanoparticles enable deep proteomics studies at scale by leveraging tunable nano–bio interactions. Proceedings of the National Academy of Sciences 119, e2106053119 (2022).

17. Donovan, M. K. R. et al. Functionally distinct BMP1 isoforms show an opposite pattern of abundance in plasma from non-small cell lung cancer subjects and controls. 2022.01.07.475393 Preprint at 10.1101/2022.01.07.475393 (2023).

18. Kverneland, A. H., Østergaard, O., Emdal, K. B., Svane, I. M. & Olsen, J. V. Differential ultracentrifugation enables deep plasma proteomics through enrichment of extracellular vesicles. PROTEOMICS n/a, 2200039.

19. Simonsen, J. B. What Are We Looking At? Extracellular Vesicles, Lipoproteins, or Both? Circulation Research 121, 920–922 (2017).

20. Iliuk, A. et al. Plasma-Derived Extracellular Vesicle Phosphoproteomics through Chemical Affinity Purification. J Proteome Res 19, 2563–2574 (2020).

21. Wu, X., Li, L., Iliuk, A. & Tao, W. A. Highly Efficient Phosphoproteome Capture and Analysis from Urinary Extracellular Vesicles. J. Proteome Res. 17, 3308–3316 (2018).

22. Hadisurya, M. et al. Quantitative proteomics and phosphoproteomics of urinary extracellular vesicles define putative diagnostic biosignatures for Parkinson’s disease. Commun Med 3, 1–19 (2023).

23. Reynolds, S. M., Käll, L., Riffle, M. E., Bilmes, J. A. & Noble, W. S. Transmembrane Topology and Signal Peptide Prediction Using Dynamic Bayesian Networks. PLoS Comput Biol 4, e1000213 (2008).

24. Hallgren, J. et al. DeepTMHMM predicts alpha and beta transmembrane proteins using deep neural networks. 2022.04.08.487609 Preprint at 10.1101/2022.04.08.487609 (2022).

25. Batth, T. S. et al. Protein Aggregation Capture on Microparticles Enables Multipurpose Proteomics Sample Preparation. Mol Cell Proteomics 18, 1027–1035 (2019).

26. Bortel, P. et al. Systematic optimization of automated phosphopeptide enrichment for high-sensitivity phosphoproteomics. Mol Cell Proteomics 100754 (2024) doi:10.1016/j.mcpro.2024.100754.

27. Koenig, C. et al. Protocol for high-throughput semi-automated label-free- or TMT-based phosphoproteome profiling. STAR Protoc 4, 102536 (2023).

28. Ishida, T. et al. Application of peptides with an affinity for phospholipid membranes during the automated purification of extracellular vesicles. Sci Rep 10, 18718 (2020).

29. Hughes, C. S. et al. Single-pot, solid-phase-enhanced sample preparation for proteomics experiments. Nature Protocols 14, 68–85 (2019).

30. Searle, B. C. et al. Chromatogram libraries improve peptide detection and quantification by data independent acquisition mass spectrometry. Nat Commun 9, 5128 (2018).

31. Pino, L. K., Just, S. C., MacCoss, M. J. & Searle, B. C. Acquiring and Analyzing Data Independent Acquisition Proteomics Experiments without Spectrum Libraries. Mol Cell Proteomics 19, 1088–1103 (2020).

32. Searle, B. C. et al. Generating high quality libraries for DIA MS with empirically corrected peptide predictions. Nat Commun 11, 1548 (2020).

33. Demichev, V., Messner, C. B., Vernardis, S. I., Lilley, K. S. & Ralser, M. DIA-NN: neural networks and interference correction enable deep proteome coverage in high throughput. Nat Methods 17, 41–44 (2020).

34. Metatla, I. et al. Neat plasma proteomics: getting the best out of the worst. Clin Proteomics 21, 22 (2024).

35. Chen, E. Y. et al. Enrichr: interactive and collaborative HTML5 gene list enrichment analysis tool. BMC Bioinformatics 14, 128 (2013).

36. Kuleshov, M. V. et al. Enrichr: a comprehensive gene set enrichment analysis web server 2016 update. Nucleic Acids Res 44, W90–97 (2016).

37. Binder, J. X. et al. COMPARTMENTS: unification and visualization of protein subcellular localization evidence. Database 2014, bau012 (2014).

38. Théry, C. et al. Minimal information for studies of extracellular vesicles 2018 (MISEV2018): a position statement of the International Society for Extracellular Vesicles and update of the MISEV2014 guidelines. Journal of Extracellular Vesicles 7, 1535750 (2018).

39. Dozio, V. & Sanchez, J.-C. Characterisation of extracellular vesicle-subsets derived from brain endothelial cells and analysis of their protein cargo modulation after TNF exposure. J Extracell Vesicles 6, 1302705 (2017).

40. Ratajczak, M. Z. & Ratajczak, J. Extracellular microvesicles/exosomes: discovery, disbelief, acceptance, and the future? Leukemia 34, 3126–3135 (2020).

41. Bağcı, C. et al. Overview of extracellular vesicle characterization techniques and introduction to combined reflectance and fluorescence confocal microscopy to distinguish extracellular vesicle subpopulations. Neurophotonics 9, 021903 (2022).

42. Jeppesen, D. K. et al. Reassessment of Exosome Composition. Cell 177, 428–445.e18 (2019).

43. Fiandaca, M. S. et al. Identification of pre-clinical Alzheimer’s disease by a profile of pathogenic proteins in neurally-derived blood exosomes: a case-control study. Alzheimers Dement 11, 600–607.e1 (2015).

44. Goetzl, E. J. et al. Altered lysosomal proteins in neural-derived plasma exosomes in preclinical Alzheimer disease. Neurology 85, 40–47 (2015).

45. Mustapic, M. et al. Plasma Extracellular Vesicles Enriched for Neuronal Origin: A Potential Window into Brain Pathologic Processes. Frontiers in Neuroscience 11, (2017).

46. Ali Moussa, H. Y., et al. Single Extracellular Vesicle Analysis Using Flow Cytometry for Neurological Disorder Biomarkers. Front Integr Neurosci 16, 879832 (2022).

47. Rai, A., Fang, H., Claridge, B., Simpson, R. J. & Greening, D. W. Proteomic dissection of large extracellular vesicle surfaceome unravels interactive surface platform. J Extracell Vesicles 10, e12164 (2021).

48. Lischnig, A., Bergqvist, M., Ochiya, T. & Lässer, C. Quantitative Proteomics Identifies Proteins Enriched in Large and Small Extracellular Vesicles. Mol Cell Proteomics 21, 100273 (2022).

49. Boersma, H. H. et al. Past, Present, and Future of Annexin A5: From Protein Discovery to Clinical Applications*. Journal of Nuclear Medicine 46, 2035–2050 (2005).

50. Vermes, I., Haanen, C., Steffens-Nakken, H. & Reutelingsperger, C. A novel assay for apoptosis. Flow cytometric detection of phosphatidylserine expression on early apoptotic cells using fluorescein labelled Annexin V. J Immunol Methods 184, 39–51 (1995).

51. Woo, H.-K. et al. Characterization and modulation of surface charges to enhance extracellular vesicle isolation in plasma. Theranostics 12, 1988–1998 (2022).

52. Ashburner, M. et al. Gene Ontology: tool for the unification of biology. Nat Genet 25, 25–29 (2000).

53. Pino, L. K. et al. Matrix-Matched Calibration Curves for Assessing Analytical Figures of Merit in Quantitative Proteomics. J. Proteome Res. 19, 1147–1153 (2020).

54. Heil, L. R., Remes, P. M. & MacCoss, M. J. Comparison of Unit Resolution Versus High-Resolution Accurate Mass for Parallel Reaction Monitoring. J Proteome Res 20, 4435–4442 (2021).

55. Pino, L. K. et al. The Skyline ecosystem: Informatics for quantitative mass spectrometry proteomics. Mass Spectrom Rev 39, 229–244 (2020).

56. Rajendran, L. et al. Alzheimer’s disease β-amyloid peptides are released in association with exosomes. Proceedings of the National Academy of Sciences 103, 11172–11177 (2006).

57. Cone, A. S. et al. Alix and Syntenin-1 direct amyloid precursor protein trafficking into extracellular vesicles. BMC Mol Cell Biol 21, 58 (2020).

58. Yuyama, K., Sun, H., Mitsutake, S. & Igarashi, Y. Sphingolipid-modulated Exosome Secretion Promotes Clearance of Amyloid-β by Microglia. J Biol Chem 287, 10977–10989 (2012).

59. Goetzl, E. J., Abner, E. L., Jicha, G. A., Kapogiannis, D. & Schwartz, J. B. Declining levels of functionally specialized synaptic proteins in plasma neuronal exosomes with progression of Alzheimer’s disease. FASEB J 32, 888–893 (2018).

60. Zhang, N., Gu, D., Meng, M. & Gordon, M. L. TDP-43 Is Elevated in Plasma Neuronal-Derived Exosomes of Patients With Alzheimer’s Disease. Front Aging Neurosci 12, 166 (2020).

61. Stuendl, A. et al. α-Synuclein in Plasma-Derived Extracellular Vesicles Is a Potential Biomarker of Parkinson’s Disease. Mov Disord 36, 2508–2518 (2021).

62. Shi, M. et al. Plasma exosomal α-synuclein is likely CNS-derived and increased in Parkinson’s disease. Acta Neuropathol 128, 639–650 (2014).

63. Chung, C.-C., Chan, L., Chen, J.-H., Hung, Y.-C. & Hong, C.-T. Plasma Extracellular Vesicle α-Synuclein Level in Patients with Parkinson’s Disease. Biomolecules 11, 744 (2021).

64. Ruan, Z. Extracellular vesicles drive tau spreading in Alzheimer’s disease. Neural Regen Res 17, 328–329 (2021).

65. Ruan, Z. et al. Alzheimer’s disease brain-derived extracellular vesicles spread tau pathology in interneurons. Brain 144, 288–309 (2021).

66. Kai, K. et al. Relationship between Eating Disturbance and Dementia Severity in Patients with Alzheimer’s Disease. PLoS One 10, e0133666 (2015).

67. Hartman, Y. A. W., Karssemeijer, E. G. A., van Diepen, L. A. M., Olde Rikkert, M. G. M. & Thijssen, D. H. J. Dementia Patients Are More Sedentary and Less Physically Active than Age- and Sex-Matched Cognitively Healthy Older Adults. Dementia and Geriatric Cognitive Disorders 46, 81–89 (2018).

68. Courtin, E. & Knapp, M. Social isolation, loneliness and health in old age: a scoping review. Health Soc Care Community 25, 799–812 (2017).

69. Plubell, D. L. et al. Can we put Humpty Dumpty back together again? What does protein quantification mean in bottom-up proteomics? bioRxiv 2021.01.25.428175 (2021) doi:10.1101/2021.01.25.428175.

70. Ando, K. et al. Dysregulation of Phosphoinositide 5-Phosphatases and Phosphoinositides in Alzheimer’s Disease. Front Neurosci 15, 614855 (2021).

71. Mostafavi, S. et al. A molecular network of the aging human brain provides insights into the pathology and cognitive decline of Alzheimer’s disease. Nat Neurosci 21, 811–819 (2018).

72. Kam, T.-I. et al. FcγRIIb-SHIP2 axis links Aβ to tau pathology by disrupting phosphoinositide metabolism in Alzheimer’s disease model. eLife 5, e18691.

73. Lee, H. N. et al. Aβ modulates actin cytoskeleton via SHIP2-mediated phosphoinositide metabolism. Sci Rep 9, 15557 (2019).

74. Lai, Y. et al. Identification of endoplasmic reticulum stress-associated genes and subtypes for prediction of Alzheimer’s disease based on interpretable machine learning. Front Pharmacol 13, 975774 (2022).

75. Chaves, M. L. et al. Serum levels of S100B and NSE proteins in Alzheimer’s disease patients. Journal of Neuroinflammation 7, 6 (2010).

76. Whitaker-Azmitia, P. M. et al. Transgenic mice overexpressing the neurotrophic factor S-100 beta show neuronal cytoskeletal and behavioral signs of altered aging processes: implications for Alzheimer’s disease and Down’s syndrome. Brain Res 776, 51–60 (1997).

77. Angelopoulou, E., Paudel, Y. N. & Piperi, C. Emerging role of S100B protein implication in Parkinson’s disease pathogenesis. Cell Mol Life Sci 78, 1445–1453 (2021).

78. Axenhus, M., Winblad, B., Tjernberg, L. O. & Schedin-Weiss, S. Huntingtin Levels are Elevated in Hippocampal Post-Mortem Samples of Alzheimer’s Disease Brain. Curr Alzheimer Res 17, 858– 867 (2020).

79. Saha, I. et al. The AAA+ chaperone VCP disaggregates Tau fibrils and generates aggregate seeds in a cellular system. Nat Commun 14, 560 (2023).

80. Kaleem, M., Zhao, A., Hamshere, M. & Myers, A. J. Identification of a novel valosin-containing protein polymorphism in late-onset Alzheimer’s disease. Neurodegener Dis 4, 376–381 (2007).

81. Ishigaki, S. et al. Physical and functional interaction between Dorfin and Valosin-containing protein that are colocalized in ubiquitylated inclusions in neurodegenerative disorders. J Biol Chem 279, 51376–51385 (2004).

82. Roux, P. P. & Blenis, J. ERK and p38 MAPK-Activated Protein Kinases: a Family of Protein Kinases with Diverse Biological Functions. Microbiol Mol Biol Rev 68, 320–344 (2004).

83. Culbert, A. A. et al. MAPK-activated protein kinase 2 deficiency in microglia inhibits pro-inflammatory mediator release and resultant neurotoxicity. Relevance to neuroinflammation in a transgenic mouse model of Alzheimer disease. J Biol Chem 281, 23658–23667 (2006).

84. Cayli, S. et al. COP9 Signalosome Interacts ATP-dependently with p97/Valosin-containing Protein (VCP) and Controls the Ubiquitination Status of Proteins Bound to p97/VCP. J Biol Chem 284, 34944–34953 (2009).

85. Cornelissen, T. et al. The deubiquitinase USP15 antagonizes Parkin-mediated mitochondrial ubiquitination and mitophagy. Hum Mol Genet 23, 5227–5242 (2014).

86. Merchant, J. P. et al. Predictive network analysis identifies JMJD6 and other potential key drivers in Alzheimer’s disease. Commun Biol 6, 1–19 (2023).

87. Dobri, A.-M., Dudău, M., Enciu, A.-M. & Hinescu, M. E. CD36 in Alzheimer’s Disease: An Overview of Molecular Mechanisms and Therapeutic Targeting. Neuroscience 453, 301–311 (2021).

88. Vapnik, V. N. The Nature of Statistical Learning Theory. (Springer, New York, NY, 2000). doi:10.1007/978-1-4757-3264-1.

89. Shahid, M. et al. Illusory Responses across the Lewy Body Disease Spectrum. Ann Neurol 93, 702–714 (2023).

90. Amodei, D. et al. Improving Precursor Selectivity in Data-Independent Acquisition Using Overlapping Windows. J Am Soc Mass Spectrom 30, 669–684 (2019).

91. Ma, Z. Q. et al. IDPicker 2.0: Improved protein assembly with high discrimination peptide identification filtering. J.Proteome.Res. 8, 3872–3881 (2009).

92. Benjamini, Y. & Hochberg, Y. Controlling the False Discovery Rate: A Practical and Powerful Approach to Multiple Testing. Journal of the Royal Statistical Society. Series B (Methodological) 57, 289–300 (1995).

