## Supplementary Materials for "Mag-Net: Rapid enrichment of membrane-bound particles enables high coverage quantitative analysis of the plasma proteome"

### SUPPLEMENTARY FIGURES

- Figure S1: Volcano plot of human plasma versus Mag-Net enriched EVs with representative markers highlighted.
- Figure S2: Volcano plot comparing the use of MagReSyn Hydroxyl versus SAX beads for protein aggregation capture (PAC) digestion.
- Figure S3: Volcano plot comparing the preparation of total plasma using PAC with Hydroxyl beads versus the use of Mag-Net membrane particle capture with Hydroxyl beads.
- Figure S4: Volcano plot comparing the difference between the use of MagResyn Hydroxyl versus SAX beads for Mag-Net.
- Figure S5: Example matrix matched calibration curves for the EV marker proteins CD9 and NCAM1 and the (V)LDL marker APOB.
- Figure S6: Assessment of plasma freeze thaw cycles on the relative peptide abundance.

- Figure S7: Selected proteins elevated in ADD that are known to be involved in vesicle mediated transport.
- Figure S8: Selected proteins elevated in ADD that are known to be involved in lipid metabolism.
- Figure S9: Selected proteins decreased in ADD that are known to be involved in ubiquitin proteosome mediated protein degradation.
- Figure S10: Individual ROC curve analysis illustrating the performance of a trained hard-margin support vector machine (SVM) with a linear kernel.

*Supplementary Figure S1*

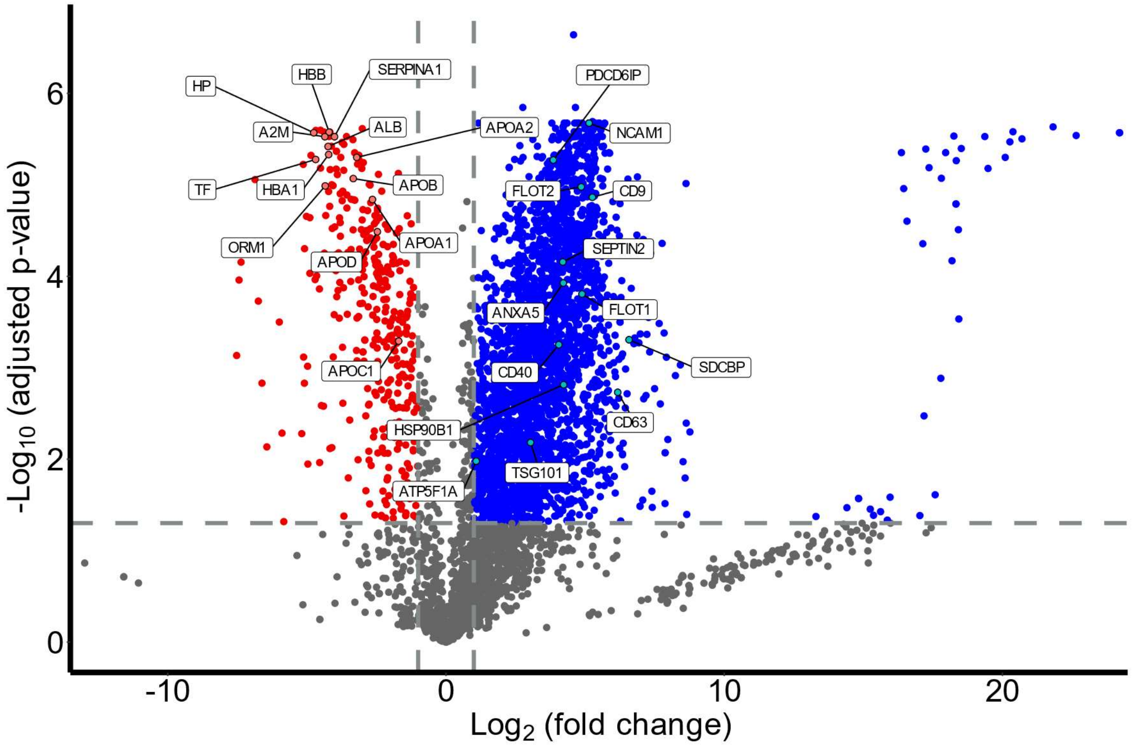


Volcano plot of the protein abundance from human plasma (red) versus plasma prepared using Mag-Net (blue). Representative proteins used as extracellular vesicle markers and classic plasma protein markers are highlighted. The figure highlights the enrichment of vesicle proteins and depletion of classic abundant plasma and lipoprotein particle proteins.

***Supplementary Figure S2***

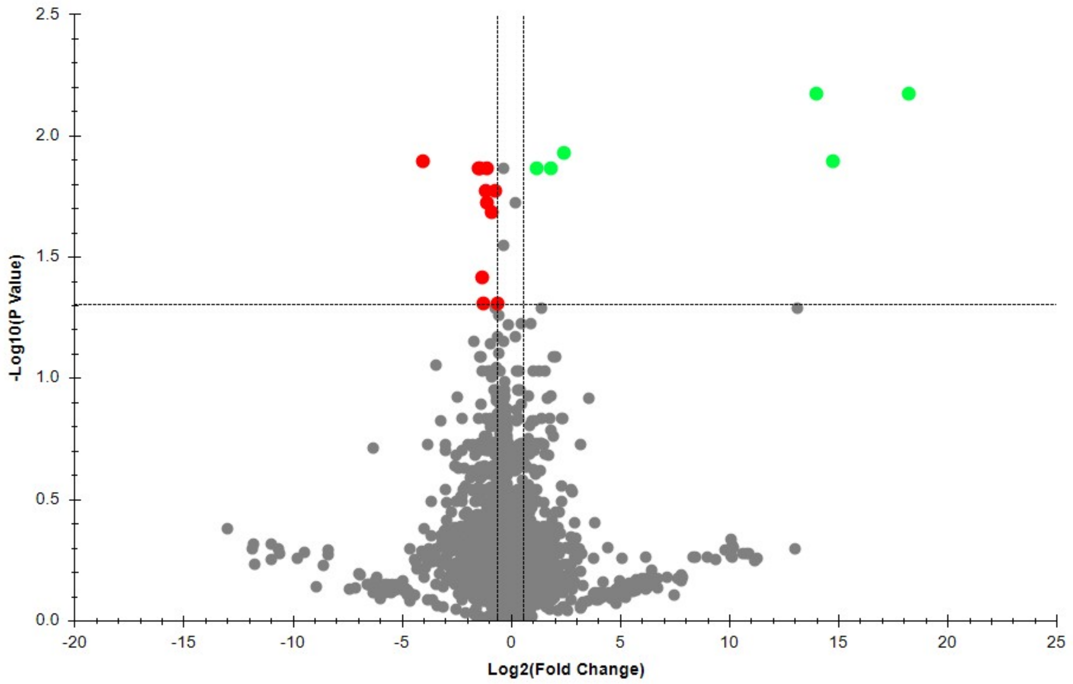


Volcano plot comparing the use of MagReSyn **Hydroxyl** vs **SAX** Beads for the digest of unfractionated plasma using protein aggregation capture (PAC) digestion. There were very few proteins with a differential abundance between the two bead types. This experiment rules out the use of SAX beads in the PAC digestion step as an explanation for the increase in proteins via Mag-Net.

***Supplementary Figure S3***

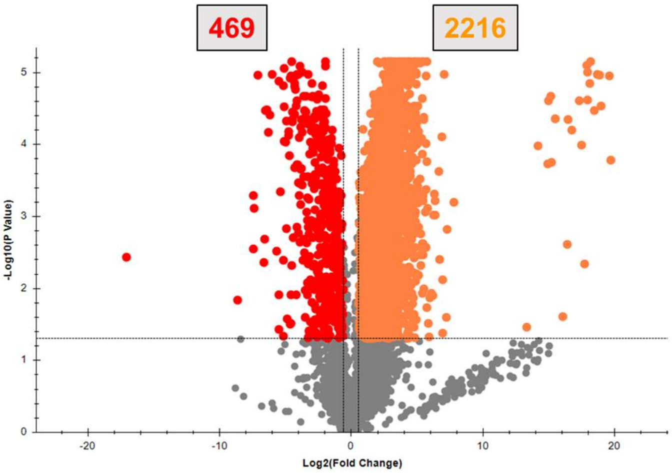


Jensen COMPARTMENTS

| Index | Name | P-value | Adjusted p-value | Odds Ratio | Combined score |
|---|---|---|---|---|---|
| 1 | membrane-bounded vesicle | 5.02E-239 | 1.03E-235 | 5.14 | 2819.43 |
| 2 | Extracellular vesicle | 5.06E-226 | 4.66E-223 | 5.41 | 2806.32 |
| 3 | Extracellular organelle | 6.80E-226 | 4.66E-223 | 5.41 | 2803.06 |
| 4 | Extracellular exosome | 6.85E-225 | 3.52E-222 | 5.4 | 2787.62 |
| 5 | Extracellular region part | 4.67E-188 | 1.92E-185 | 4.23 | 1823.67 |
| 6 | Extracellular region | 2.09E-159 | 7.17E-157 | 3.64 | 1328.37 |
| 7 | Platelet alpha granule | 1.47E-25 | 8.86E-24 | 13.65 | 780.31 |
| 8 | cell-cell adherens junction | 1.20E-44 | 1.89E-42 | 5.89 | 595.43 |
| 9 | Integrin alphaIIb-beta3 complex | 2.30E-27 | 1.63E-25 | 8.82 | 540.94 |
| 10 | Platelet alpha granule lumen | 1.15E-18 | 4.20E-17 | 12.8 | 528.78 |

Differential analysis between **total (unfractionated) Plasma PAC digest using Hydroxyl** vs **the Mag-Net protocol using Hydroxyl** Beads. These data illustrate that a hyperporous MagReSyn without the charge from the quaternary ammonium functionality, will still capture and enrich membrane particles.

### *Supplementary Figure S4*

GO Cellular Component

| Index | Name | P-value | Adjusted p-value | Odds Ratio | Combined score |
|---|---|---|---|---|---|
| 1 | collagen-containing extracellular matrix (GO:0062023) | 7.47E-29 | 1.38E-26 | 9.83 | 636.83 |
| 2 | endoplasmic reticulum lumen (GO:0005788) | 1.59E-15 | 1.20E-13 | 7.36 | 250.93 |
| 3 | high-density lipoprotein particle (GO:0034364) | 1.95E-15 | 1.20E-13 | 82.13 | 2781.8 |
| 4 | spherical high-density lipoprotein particle (GO:0034366) | 3.04E-12 | 1.41E-10 | 413.25 | 10958.82 |
| 5 | membrane attack complex (GO:0005579) | 2.31E-11 | 8.20E-10 | 117960 | 2888975.47 |
| 6 | triglyceride-rich plasma lipoprotein particle (GO:0034385) | 3.73E-11 | 8.20E-10 | 67.65 | 1624.52 |
| 7 | very-low-density lipoprotein particle (GO:0034361) | 3.73E-11 | 8.20E-10 | 67.65 | 1624.52 |
| 8 | secretory granule lumen (GO:0034774) | 3.73E-11 | 8.20E-10 | 5.53 | 132.8 |
| 9 | intracellular organelle lumen (GO:0070013) | 4.40E-11 | 8.20E-10 | 3.49 | 83.14 |
| 10 | chylomicron (GO:0042627) | 4.43E-11 | 8.20E-10 | 137.74 | 3283.61 |

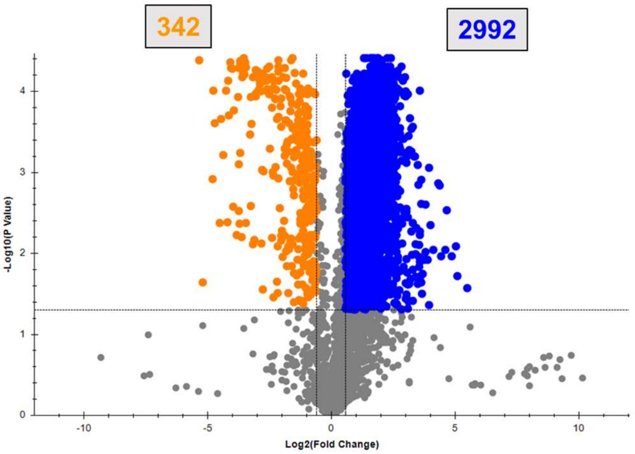


GO Cellular Component

| Index | Name | P-value | Adjusted p-value | Odds Ratio | Combined score |
|---|---|---|---|---|---|
| 1 | intracellular organelle lumen (GO:0070013) | 4.65E-57 | 1.98E-54 | 3.54 | 459.38 |
| 2 | focal adhesion (GO:0005925) | 1.69E-55 | 2.59E-53 | 5.61 | 707.18 |
| 3 | cell-substrate junction (GO:0030055) | 1.83E-55 | 2.59E-53 | 5.53 | 697.17 |
| 4 | secretory granule lumen (GO:0034774) | 3.27E-54 | 3.48E-52 | 6.44 | 792.66 |
| 5 | bounding membrane of organelle (GO:0098588) | 5.73E-46 | 4.88E-44 | 3.3 | 343.58 |
| 6 | mitochondrial membrane (GO:0031966) | 1.25E-41 | 8.91E-40 | 4.03 | 379.7 |
| 7 | ficolin-1-rich granule (GO:0101002) | 1.07E-35 | 6.52E-34 | 6.97 | 561.43 |
| 8 | secretory granule membrane (GO:0030667) | 1.40E-35 | 7.44E-34 | 5.09 | 408.4 |
| 9 | mitochondrial matrix (GO:0005759) | 2.72E-35 | 1.29E-33 | 4.32 | 344.15 |
| 10 | endosome membrane (GO:0010008) | 3.27E-32 | 1.40E-30 | 4.24 | 307.46 |

Differential analysis of proteins enriched using Mag-Net with either **Hydroxyl** vs **SAX** Beads. Because both hydroxyl and SAX beads are both capable of enriching membrane particles, we compared the proteins that were selectively enriched or depleted relative to each other. From these data, the SAX beads are more specific to EVs and are depleted in lipoprotein and chylomicron particles relative to the hydroxyl beads.

*Supplementary Figure S5*

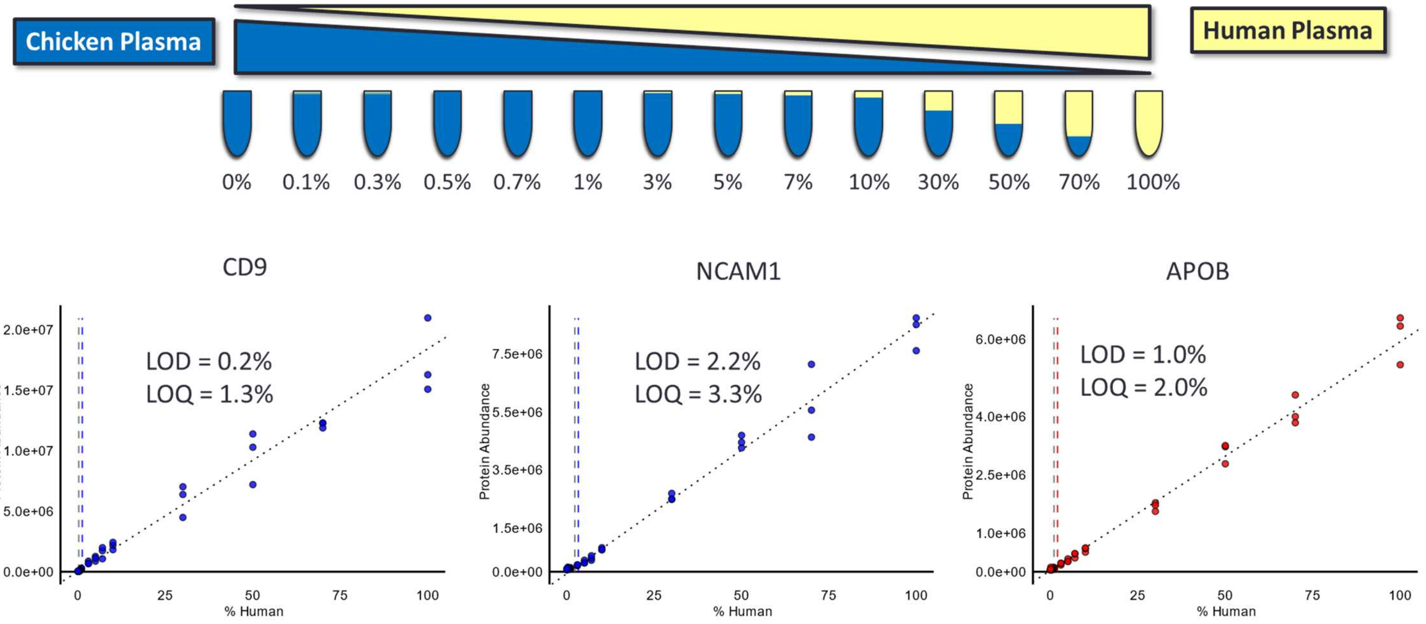


We assessed the quantitative accuracy and linearity using matrix match calibration curves. We took a human plasma sample and diluted it volumetrically with chicken plasma spanning a range of $10^3$. There separate and independent calibration curves were produced. All samples (14 x 3 = 42 total) were prepared individually and measured. Shown are individual calibration curves for the EV markers CD9 and NCAM1. Also shown is the low-density lipoprotein marker APOB. Despite APOB being depleted using Mag-Net, most depleted proteins still have a quantitative linear response. Full Skyline document for all peptides and proteins is available on Panorama Public.

***Supplementary Figure S6***

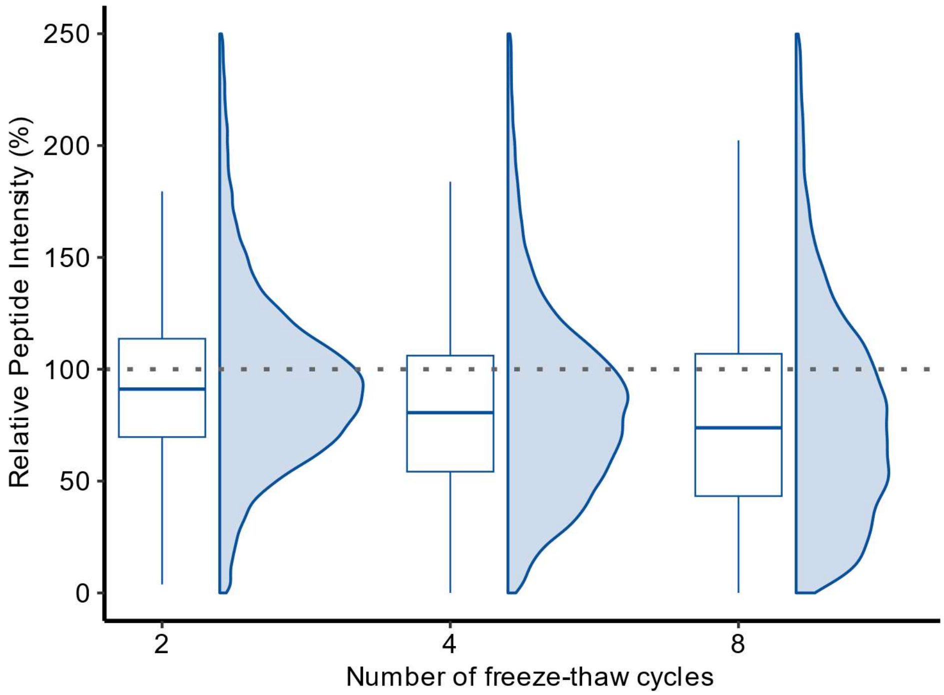


Assessment of plasma freeze thaw cycles on the relative abundance of peptides using Mag-Net. A single plasma sample was aliquoted. Aliquots were collected from 1, 2, 4, and 8 freeze thaw cycles. The aliquots of the same plasma sample but from different freeze thaw cycles were randomized, prepared and measured together. The abundance of each peptide was then measured relative to the respective peptide abundance obtained from a single freeze thaw cycle. The median relative peptide abundance was >75% even with 8 plasma freeze thaw cycles.

### *Supplementary Figure S7*

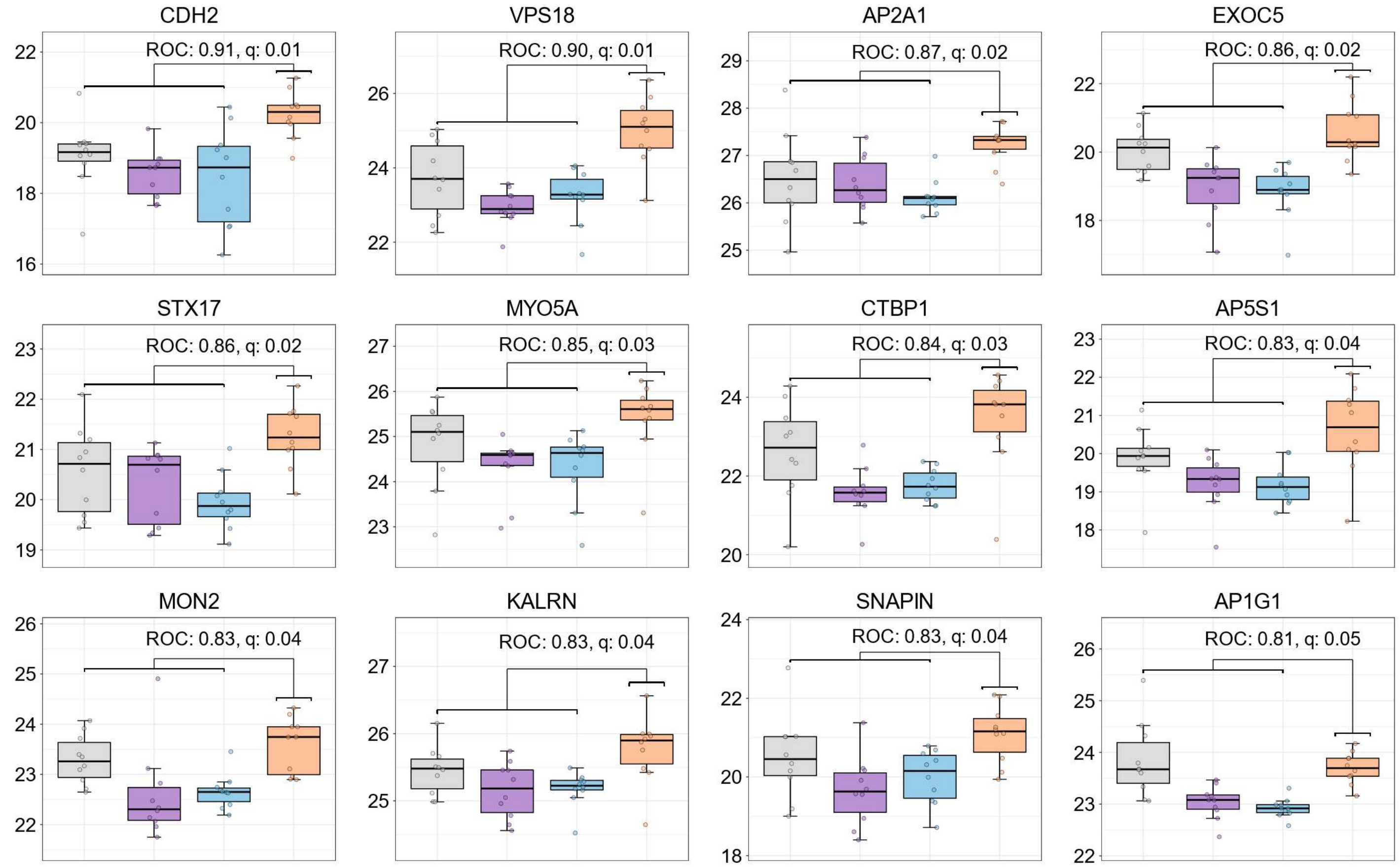


Selected proteins elevated in ADD that are known to be involved in vesicle mediated transport. The grey = HCN, purple = PDCN, blue = PDD, and orange = ADD.

*Supplementary Figure S8*

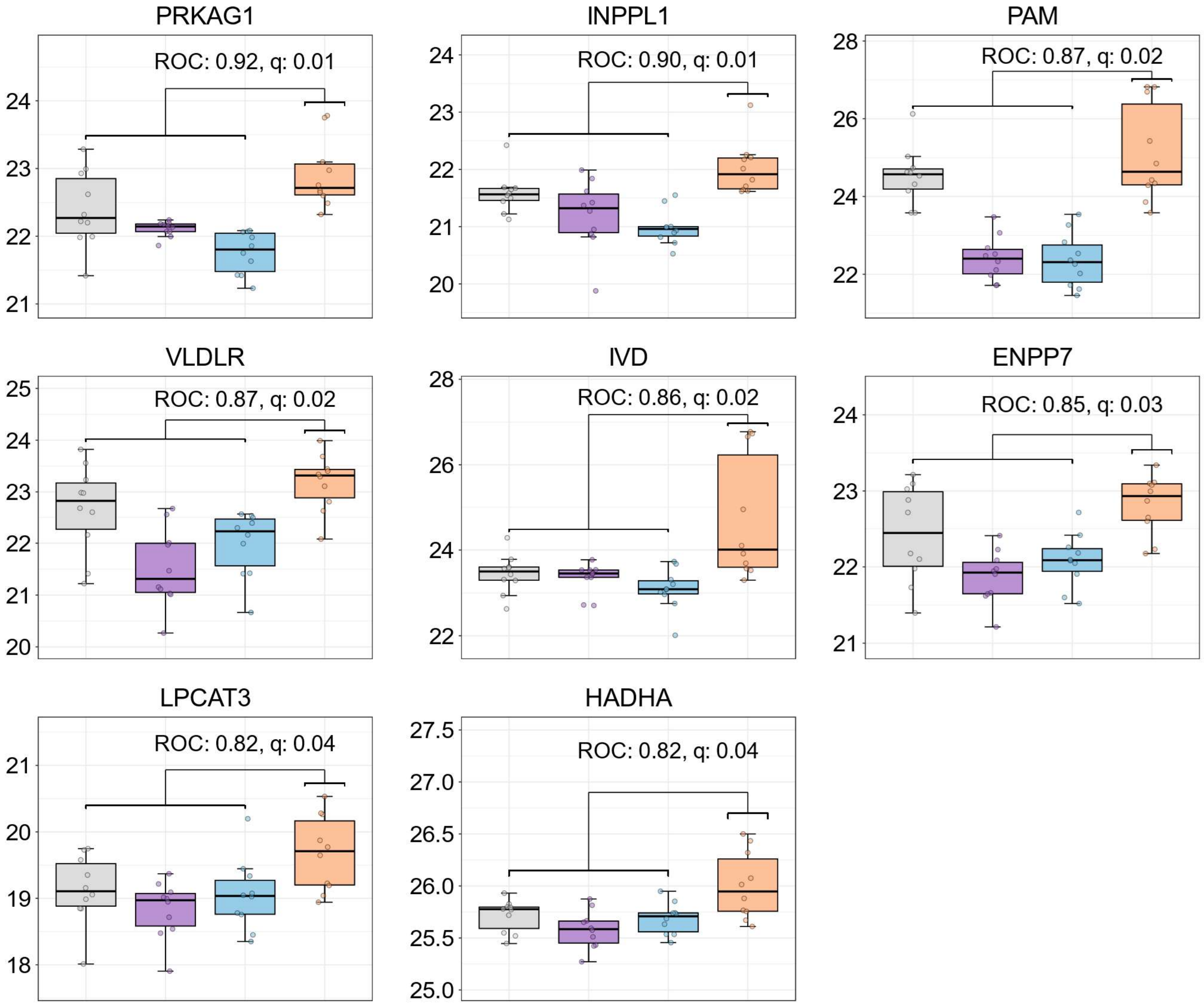


Selected proteins elevated in ADD that are known to be involved in lipid metabolism. The grey = HCN, purple = PDCN, blue = PDD, and orange = ADD.

***Supplementary Figure S9***

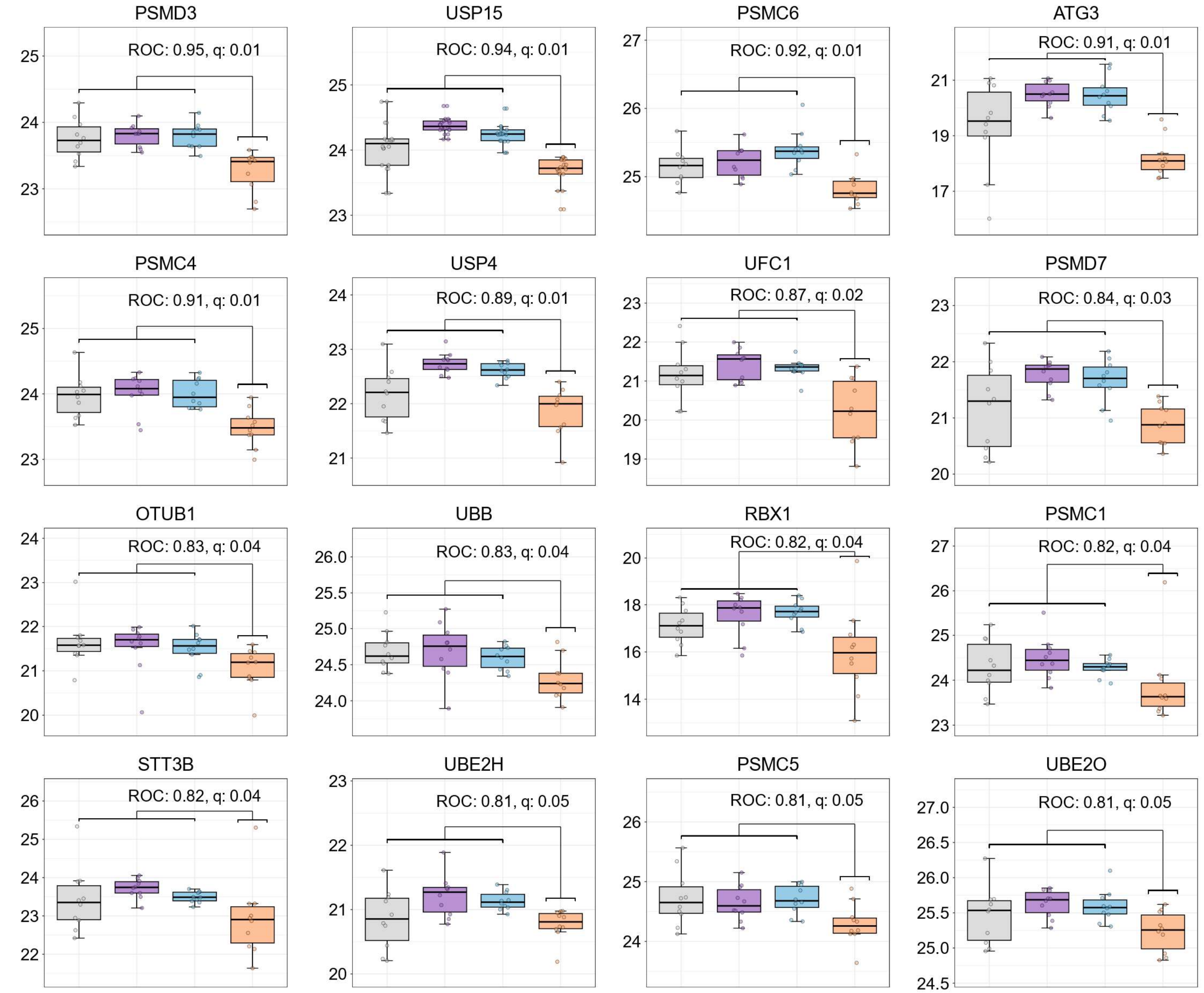


Selected proteins decreased in ADD that are known to be involved in ubiquitin proteosome mediated protein degradation. The grey = HCN, purple = PDCN, blue = PDD, and orange = ADD.

*Supplementary Figure S10*

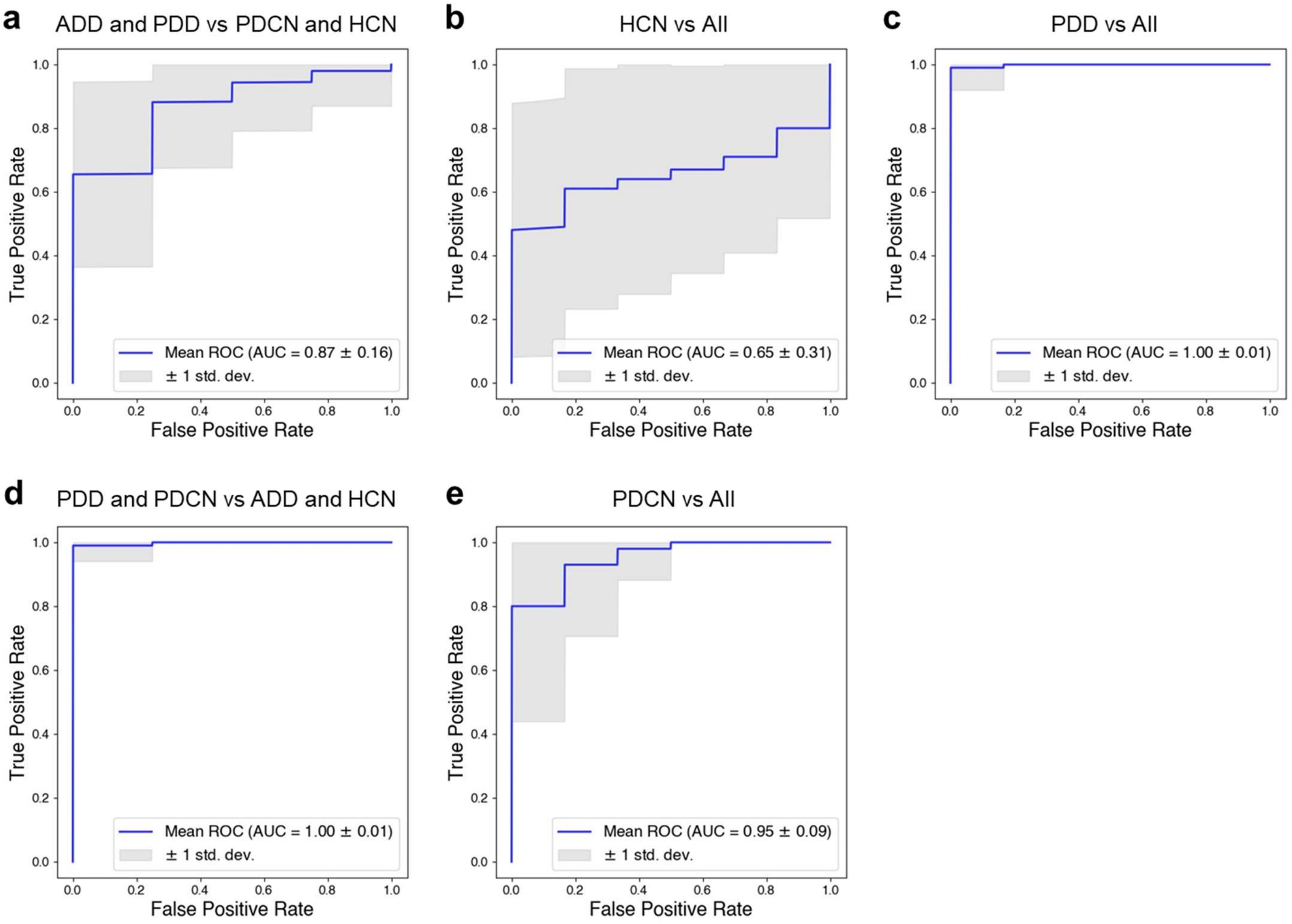


We developed an SVM classifier to perform six pair wise analyses. Figure a shows an ROC curve analysis for dementia (ADD and PDD) versus cognitively normal (HCN and PDCN). Figure b shows HCN versus all others. Figure c shows PDD versus all others. Figure d shows Parkinson's disease (PDCN and PDD) versus non-Parkinson's disease (ADD and HCN). Figure e shows PDCN versus all others. The final ROC analysis of ADD versus all others is in Figure 7b of the main text.
